# Microbiota impact *Drosophila* ageing via *Acetobacter, Tachykinin,* and *TkR99D*

**DOI:** 10.1101/2025.07.31.667994

**Authors:** Diana Marcu, David R Sannino, Anthony J Dornan, Rita Ibrahim, Atharv Kapoor, Miriam Wood, Adam J Dobson

**Affiliations:** School of Molecular Biosciences, College of Medical Veterinary and Life Sciences, University of Glasgow, G12 8QQ, United Kingdom; Cellular Analysis Facility - Shared Research Facilities, College of Medical Veterinary and Life Sciences, University of Glasgow, G12 8TA, United Kingdom

## Abstract

Gut microbiota exert an evolutionarily conserved influence on ageing, from invertebrates to humans. How do microbes that are physically confined to the gut lumen affect the systemic physiological process of ageing? In female *Drosophila*, we show that microbiota increase expression of the peptide hormone *Tachykinin (Tk)*, which corresponds to reduced lifespan. *Tk* is required for microbiota to shorten lifespan, with knockdown rendering flies constitutively long-lived even in the presence of an intact microbiota. This lifespan extension does not come with canonical costs to fecundity or feeding, but impacts on triacylglyceride (TAG) storage suggest adaptive functions in metabolic homeostasis. In flies with defined (gnotobiotic) microbiotas, we show that we can model *Tk*-dependent effects of microbiota on lifespan and TAG by monoassociation with *Acetobacter pomorum*. These effects require *Tk* in the midgut, and the cognate TK receptor *TkR99D* in neurons, implicating a microbiota-gut-neuron relay. This relay also appears to compromise gut barrier function in aged flies, indicating roles in healthspan as well as lifespan. However, the effect of *TkR99D* is independent of its reported role in insulin signalling and adipokinetic hormone signalling which, respectively, are canonical regulators of lifespan and TAG metabolism, suggesting a non-canonical role for *TkR99D* elsewhere in the nervous system. Altogether our results implicate a microbiota-gut-neuron axis in ageing, via a specific bacterium modulating activity of a specific and evolutionarily-conserved hormone.

## Introduction

Gut microbiota influence lifespan and ageing across species. This effect appears to be evolutionarily conserved from the invertebrate models *Caenorhabditis elegans* and *Drosophila melanogaster* (Brummel et al., 2004; Cabreiro et al., 2013; Houthoofd et al., 2002; Matthews et al., 2020; Obata et al., 2018; Pryor et al., 2019; Sannino et al., 2018), through to vertebrates including fish and rodents (Gordon et al., 1966; Smith et al., 2017; Snyder et al., 1990; Tazume et al., 1991; Valenzano et al., 2015). Microbiota are also thought to play a role in human ageing (Johansen et al., 2023; OToole and Jeffery, 2015), supported by mendelian randomisation analysis indicating a putatively causal role (Liu et al., 2023) (but see also (Gagnon et al., 2023)). Gut microbiota are physically confined to the gut lumen, yet they influence ageing at an organismal level. By what mechanisms do microbiota modulate these systemic processes?

Evolutionary theories of ageing lead us to expect that mechanisms of ageing should have adaptive functions early in life, i.e. processes under positive selection in early life have deleterious pleiotropic effects in later life, which manifest as ageing (Williams, 1957). Microbiota conform to this expectation, with their impacts on ageing appearing to trade off against widespread benefits to host health early in life. For example, microbiota commonly promote host development time, even with the same species of bacteria demonstrated to benefit host development across different phyla of animals (Schwarzer et al., 2016; Storelli et al., 2011; A. Wong et al., 2014). This motivates research to characterise how to decouple beneficial effects of microbiota early in life from deleterious effects in late life.

The capacity of microbiota to impact systemic processes, from a physically confined niche in the intestinal lumen, implicates endocrine signalling between organs. A role mediated by endocrine signalling is also consistent with the pleiotropic effects of microbiota, since hormones potently impact multiple traits. The relationship between microbiota and endocrine signalling may be bidirectional, with the microbiota altering how the gut - the biggest endocrine organ in animals - communicates lumen conditions to distal tissues, while distal tissues may direct how the gut manages the microbiota. Signalling in either direction has the potential to modulate ageing, with hormones serving as a key relay.

We chose to investigate these processes in the fruitfly *Drosophila melanogaster*. Flies can be reared axenic (i.e. without exogenous bacteria), and have a simple microbiota dominated by two families of bacteria (*Acetobacteraceae* and *Lactobacillaceae*). These bacteria can be cultured easily, and reintroduced to axenic fly eggs to generate gnotobiotic animals, enabling mapping of host phenotype to bacterial species. Fly genetics also enable tissue- and cell-type-specific genetic manipulations, e.g. expressing RNAi in specific locations, ideal for studying tissue-tissue signalling. We capitalised on these strengths to identify how defined bacteria modulate hormone signalling, and how ageing phenotypes are modulated by tissue-specific lesions in this signalling.

## Results

### A role for *Tk* in lifespan regulation by microbiota

First, we looked for evidence that microbiota modulate fly endocrine signalling. Hormone activity both to and from the gut could be relevant to impact of microbiota; but since the gut is the site of physical contact with gut microbiota, and also the largest endocrine organ, in which numbers of enteroendocrine cells (EEs) can vary plastically in adults, we reasoned that EE counts could serve as a proxy for microbial impacts of overall endocrine signalling capacity, even though other endocrine tissues could also be affected. When we counted absolute numbers of EEs in whole guts (Figure 1A), we found more cells in guts of conventionally-reared (CR) flies than in axenic flies, suggesting that microbiota enhance capacity for endocrine signalling. Previous studies have shown that microbiota reduce the percentage of EEs (Broderick et al., 2014; Liu et al., 2022), which we suggest may have been driven by an even greater microbial promotion of other epithelial cell populations, reducing proportion of EEs despite increased total numbers of EEs.

**Figure 1.**
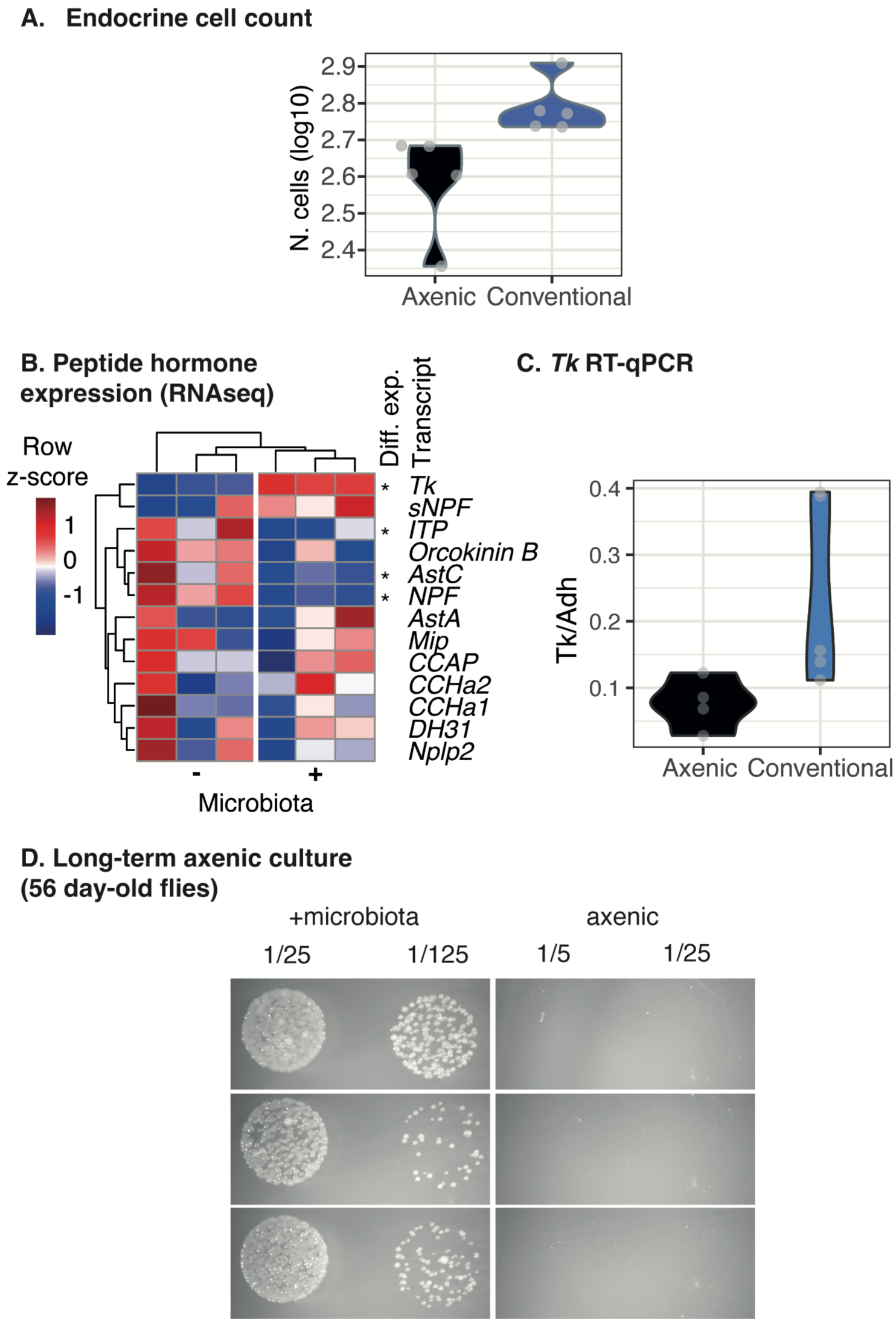
Basis for study: evidence that *Tk* responds to microbiota. **A.** Microbiota increase total number of enteroendocrine cells (EEs) in fly midgut. Number of GFP+ cells were scored in 5-day old *UAS-mCD8::GFP; Voila-Gal4* females. Wilcoxon test, W=25, P=0.008. **B.** Microbiota alter expression of peptide hormones in the *Drosophila* intestine. Transcriptome data from (Bost et al., 2018) were reanalysed. Expression of neuropeptide hormone gene transcripts are plotted, dendrograms show hierarchical clustering on Euclidian distance. Differentially expressed transcripts (DESeq2, FDR<0.1) are marked with an asterisk to right. Presence of microbiota increased expression of *Tachykinin* (top row). Software did not quantify *Burs* expression, so complementary RT-qPCR data were collected, showing no impact of microbiota (Figure S1). **C.** Confirmation that microbiota promote *Tk* expression. Expression was quantified by RT-qPCR (using cDNA generated from midgut to complement RNA-seq data). Wilcoxon test W=1, p=0.04. **D.** Confirmation of lifelong axenic culture without antibiotics. Photographs show bacterial growth (or lack of) in homogenates of pools of 3 females, reared to day 56 of adulthood either with bacteria or axenic, serial dilutions indicated at top.

EEs express 14 peptide hormones (Hung et al., 2020). We asked whether microbiota promote expression of any of these specifically, by reanalysing public transcriptome data (Bost et al., 2018) (Figure 1B). We were able to quantify expression of 13 genes, and we analysed the fourteenth (*Burs*) by RT-qPCR (Figure S1). Four hormone genes were differentially expressed. Of these four, Tachykinin (*Tk*) - which encodes orthologues of the mammalian neuropeptides Substance P and Neurokinins A/B - appeared a particularly interesting candidate, because published data indicated that *Tk* knockdown flies phenocopy the longer lifespan and metabolic phenotypes observed in axenic flies (Dobson et al., 2015; Rewitz et al., 2024; Sannino et al., 2018; Song et al., 2014; A. C.-N. Wong et al., 2014). Since the differential expression analysis was performed on public data, we validated the impact of microbes on *Tk* expression in our lab stock (*Dahomey*) by RT-qPCR (Figure 1C). This correlation between microbial promotion of *Tk* expression and shortening of lifespan, coupled to phenocopying between axenic and *Tk* knockdown flies, led us to predict that *Tk* knockdowns would be constitutively long-lived and their lifespan should no longer respond to microbiota.

We tested our prediction by assessing impact of *Tk* knockdown in flies reared with/without a microbiota. We reared flies under axenic conditions from embryo until death, to avoid potential off-target effects on the fly of antibiotic feeding (Chatzispyrou et al., 2015; Moullan et al., 2015; Ronayne et al., 2023), after confirming that we could maintain sterile culture conditions throughout the lifespan (Figure 1D). To establish the genetic role of *Tk*, we first used a ubiquitous knockdown, to avoid in the first instance the possibility that targeting one specific tissue may lead to compensatory expression in another. We drove RNAi expression using an inducible ubiquitous driver (*Daughterless-GeneSwitch* a.k.a. *DaGS)*, which we activated in 3-day-old adult females by feeding a chemical inducer, RU486 (henceforth, RU), thereby avoiding any possible confounding impacts of knockdown during development. In all experiments, we used female flies and a fully-factorial design.

Microbiota shortened lifespan relative to axenic controls (Figure 2A). However, *Tk^RNAi^* blocked this effect, rendering CR flies as long-lived as axenics. Cox Proportional Hazards analysis (CoxPH) revealed a significant *Tk^RNAi^*:microbiota interaction (p=0.05). To visualise specifically how the experimental conditions differed from one another, we used estimated marginal means (EMMs) to calculate *post hoc* differences between conditions, and to plot *post hoc* metrics of differences in survival (multiplied by −1 such that higher values corresponded to extended lifespan). This analysis revealed a significant shortening of lifespan by microbiota in CR flies relative to axenics, but *Tk^RNAi^* blocked this effect (Figure 2B), i.e. *Tk^RNAi^* extended lifespan in conventional flies, but not in axenic flies. To confirm the efficacy of *Tk* knockdown in conventional flies with an independent tool, we expressed a distinct RNAi in conventional flies, which also extended lifespan (Figure S3); and an independent recent study (Ahrentløv et al., 2025) also corroborates our findings. CR lifespan was not extended by RU in the absence of the *UAS-Tk^RNAi^* transgene (Figure S4), indicating that the *Tk:*microbiota interaction was attributable to knockdown, not off-target effects of RU feeding; and bacterial load was not altered by *Tk^RNAi^* induction, suggesting that the lifespan difference is not due to altered density of microbiota (Figure S5). Together, these results revealed that gut-microbial modulation of lifespan requires *Tk*.

**Figure 2.**
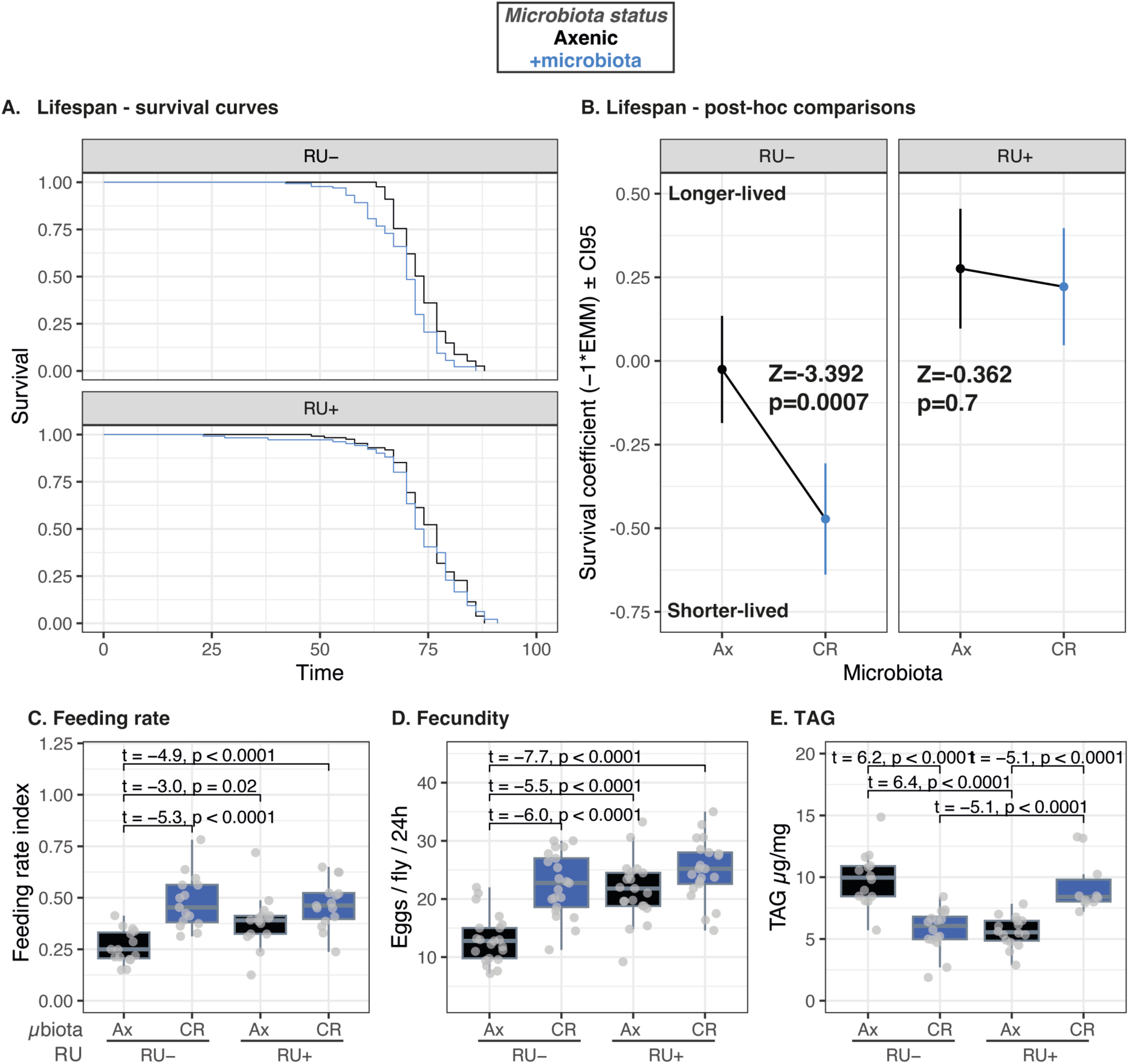
Ubiquitous *Tk* is required for microbiota to shorten lifespan and reduce TAG storage. **A.** Kaplan-Meier plots showing survival of axenic and conventionally-reared (CR) *DaGS;UAS-Tk^RNAi^* flies, faceted by absence/presence (top/bottom) of transgene activation by RU486 (RU). Cox Proportional Hazards analysis revealed an RU:microbiota interaction (p=0.05). Axenic RU- n=135, axenic RU+ n=136, CR RU- n=135, CR RU+ n=135. **B.** Post-hoc analysis of survival data shown in A. Facets show estimated marginal means of Cox Proportional Hazards model (EMMs±95% confidence intervals (CI95)), multiplied −1*n to make higher values correspond to longer lifespan. Pairwise tests (text on facets) revealed impact of microbiota only in absence of RU. **C-D.** *Tk^RNAi^* (RU+) does not block effect of microbiota on feeding rate (C) or fecundity (D). Feeding quantified as average proboscis extension rate per vial (n=5 mated females/vial, 10 days old after 7 days feeding on RU/vehicle.). Fecundity quantified as n. eggs laid per fly over 24h (n=5 females/vial). **E.** *Tk^RNAi^* (RU+) blocks reduction of TAG by microbiota. Points give TAG µg mg^−1^ in pools of five flies. The result was replicated with an independent RNAi construct (Figure S3). Panels A-E generated with RNAi construct V330743. Panels A-B, n=135-136 flies per condition. Panels C-E, Statistics on brackets are from *post-hoc* EMM analyses of linear models (C, logit-transformed data), when significant differences were detected.

### Evidence for pleiotropic roles of *Tk* between lifespan and metabolism, but not canonical tradeoffs with ageing

Tachykinins are highly pleiotropic peptides (Nässel, 2025), and interventions that extend lifespan are commonly associated with pleiotropic trade-offs early in life. For example, lifespan is expected to anticorrelate reproduction in early life (Partridge et al., 2005), while feeding behaviour, naturally selected to optimise growth and reproduction, is expected to be deleterious for ageing (Flatt, 2009; Good and Tatar, 2001; Grandison et al., 2009; Mair et al., 2005). We therefore hypothesised that *Tk^RNAi^* might block beneficial effects of microbiota on reproduction or feeding in early life. We measured steady-state feeding behaviour (proboscis extension) and a point measure of fecundity (egg laying over 24h), after one week of *Tk^RNAi^* induction in both axenic and CR flies. As expected, microbiota did indeed increase feeding and egg laying in CR flies relative to axenics (Figure 2C-D), but *Tk^RNAi^* did not reduce either fecundity or feeding. Thus, longevity in CR *Tk^RNAi^* flies appears not to be associated with canonical costs.

We looked for alternative pleiotropic traits. Microbiota have evolutionarily-conserved effects on host metabolism, with increased lipid storage observed in axenic hosts (Dobson et al., 2015; Turnbaugh et al., 2006; Vijay-Kumar et al., 2010). Since *Tk* is also implicated in mobilising fly triglyceride (TAG) stores (Song et al., 2014), we anticipated a potential tradeoff with lifespan. Without *Tk^RNAi^*, microbiota reduced TAG storage as expected, but *Tk^RNAi^* blocked this effect in two replicate experiments with distinct *Tk^RNAi^* constructs (Figure 2E, Figure S6). These results suggested a pleiotropic role for *Tk* in managing metabolic responses to microbiota in early life, with deleterious impacts for lifespan.

### *Acetobacter pomorum,* but not *Levilactobacillus brevis*, interacts with *Tk* to determine lifespan and TAG

Can the interaction between microbiota and *Tk* be attributed to particular gut bacteria? The fly microbiota, while relatively simple, still contains numerous taxa, which vary among strains and environments, but tend to contain families *Acetobacteraceae and Lactobacillaceae* (Brown et al., 2023; Chandler et al., 2011; Staubach et al., 2012; Wang and Staubach, 2018; Wong et al., 2013). We characterised the fly microbiota in our lab background by 16S rRNA amplicon sequencing, and identified amplicon sequencing variants (ASVs). Most ASVs were assigned to the family *Acetobacteraceae* (Figure 3a), and *Lactobacillaceae* were also present (among other families). *Acetobacter pomorum* and *Levilactobacillus brevis* are two well-studied species that can be considered representatives of their respective genera, on the basis of genome content (Newell et al., 2014) and phenotypic impacts (Newell and Douglas, 2014). We asked whether either *A. pomorum* or *L. brevis* alone recapitulated impacts of a CR microbiota on lifespan and TAG, and whether they interacted equivalently with *Tk^RNAi^*.

**Figure 3.**
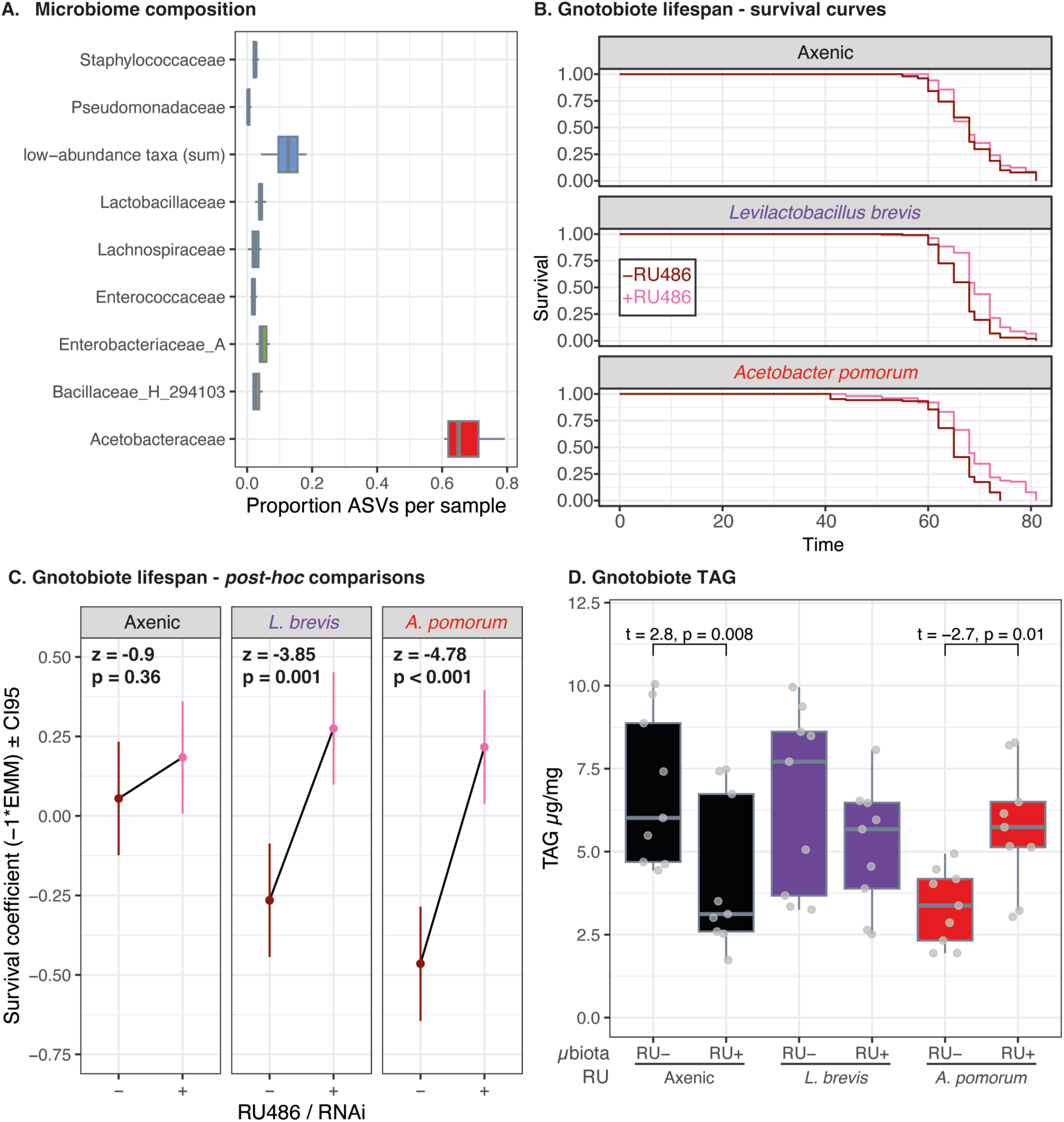
Phenotypic impact of a complete microbiota and interaction with ubiquitous *Tk^RNAi^* is recapitulated fully by *Acetobacter pomorum*, but only partially by *Levilactobacillus brevis.* **A.** 16s rRNA amplicon sequencing reveals microbiota dominated by family *Acetobacteraceae* in the *Dahomey* fly background. Plot shows proportion ASVs per sample assigned to given families. “low abundance taxa” denotes a bin of all ASVs not assigned to one of the given families. ∼170k reads per sample, n=10 samples. **B.** Kaplan-Meier plots showing survival of *DaGS;UAS-Tk^RNAi^* flies (RNAi construct V330743), cultured either under axenic or gnotobiotic conditions. Gnotobiotic flies were cultured in monoassociation with either *Acetobacter pomorum* (*Ap-*flies) or *Levilactobacillus brevis* (*Lb-*flies). Plots are faceted by microbial condition. Cox Proportional Hazards analysis revealed an RU:microbiota interaction (p=0.02). Ap RU- n=105, Ap RU+ n=105, Ax RU- n=105, Ax RU+ n=104, Lb RU- n=105, Lb RU+ n=105. **C.** Post-hoc analysis of survival curves shown in B. Plots show a survival coefficient, calculated from estimated marginal means of Cox Proportional Hazards model (EMMs±95% confidence intervals (CI95)), multiplied −1*n such that higher values correspond to longer lifespan. Pairwise tests (text on facets) revealed effect of *Tk^RNAi^* induction (+RU) was largest in *Ap-*flies, intermediate in *Lb*-flies, and no effect was observed in axenics. D. *A. pomorum,* but not *L. brevis,* recapitulates effects of a complete microbiota on TAG. Points give TAG µg mg^−1^ in pools of five flies (n=9). Significant interaction of microbiota and *TkRNAi* (RU) was detected (ANOVA F=6.96, df=2,48, p=0.002). Statistics on brackets are from *post-hoc* EMM analyses of linear models, when significant differences were detected.

We generated gnotobiotic flies, monoassociated with either *A. pomorum* (DmCS004) or *L. brevis* (DmCS003), along with axenic controls, and tested whether lifespan impacts were blocked by ubiquitous *Tk^RNAi^* (Figure 3B-C). Without *Tk^RNAi^*, axenic flies were longest-lived, and both bacteria shortened lifespan. flies associated with *A. pomorum* (henceforth *Ap*-flies) were shortest-lived, and flies associated with *L. brevis* (*Lb-*flies) were intermediate. However, *Tk^RNAi^* induction rendered flies constitutively long-lived, independent of microbiota association and equivalent to axenic controls. Thus, *Tk^RNAi^* blocked variation in lifespan caused by synthetic variation in the microbiota, and this effect was strongest in *Ap*-flies.

We quantified *Tk* expression by RT-qPCR among axenic flies, *Lb*-flies and *Ap*-flies, and asked whether change in expression upon knockdown (*DaGS, UAS-Tk^RNAi^*) correlated lifespan outcome (Figure S7). *Tk* expression was lowest in axenics, highest in *Ap*-flies, and intermediate in *Lb*-flies (Figure S7A), but RU feeding arrested *Tk* expression across all three microbiota conditions - confirming universally efficient knockdown - and this expression level was equivalent to control axenic flies (Figure S7A). Plotting *Tk* expression against statistical coefficients (EMMs) of the corresponding survival data, revealed a correlation across conditions between *Tk* expression values and lifespan (Figure S7B). Thus, microbial modulation of lifespan appears to correspond to organismal expression of *Tk*.

We then asked whether specific bacteria and *Tk* interactively modulated TAG levels in youth (Figure 3E). We found reduced TAG in *Ap*-flies relative to axenic flies, and this effect was reversed by *Tk^RNAi^*, consistent with the pleiotropic tradeoff we had observed in CR flies. However no interaction was apparent between *L. brevis* and *Tk^RNAi^*. Thus, *A. pomorum* was sufficient to modulate TAG in interaction with *Tk*, but *L. brevis* was not. We also note that in both this experiment and the equivalents in CR flies (Figure 2E, Figure S6), feeding RU to axenic flies reduced TAG. Since *Tk^RNAi^* does not reduce *Tk* expression in axenics (Figure S7), we interpret this as a possible off-target effect of RU, however such an effect cannot account for the lack of difference between axenics without RU and *Ap-*flies/*Lb-*flies with RU.

Altogether, these data demonstrated that *A. pomorum* is sufficient to fully recapitulate the impacts of a complete microbiota on both lifespan and TAG, and interacts equivalently with ubiquitous *Tk^RNAi^*. By comparison the intermediate effects in *Lb-*flies argues against *Lactobacillaceae* playing a leading role in the *Tk*-dependent regulation of lifespan and TAG.

### *A. pomorum* modulates *Tk* expression in midgut but not CNS

Our initial experiments employed ubiquitous *Tk^RNAi^*. From which specific tissue does *Tk* knockdown mediate effects of the microbiota? Plotting steady-state *Tk* expression from the FlyAtlas2 database (Leader et al., 2018) (i.e. from CR flies), revealed that *Tk* is enriched in neuronal and midgut tissues (Figure 4A). We therefore characterised metrics of Tachykinin activity from these two tissues in axenic flies, *Lb-*flies and *Ap*-flies. We examined *Tk* expression by RT-qPCR of RNA from heads and on dissected midguts. *A. pomorum* increased *Tk* expression in midgut, but not in heads; while *L. brevis* did not impact *Tk* expression in either tissue (Figure 4B). We used reporters to confirm intestinal modulation of *Tk* expression. *Acetobacter* are thought to promote Tk signalling epigenetically, by promoting histone lysine acetylation in *Tk+* cells (Jugder et al., 2021). We confirmed that lysine acetylation was labile in *Tk+* EEs by feeding axenic flies with a histone deacetylase inhibitor, trichostatin-A, and quantifying staining in *Tk+* cells in both anterior and posterior regions of the midgut (Figure 4C-D). We then examined effects of microbiota, finding increased acetyl-lysine in *Tk+* cells of *Ap-*flies, but not *Lb-*flies (Figure 4D). We asked whether this increase corresponded to elevated *Tk* expression, and indeed *Tk* reporter expression (*Tk-T2A-Gal4>UAS-*GFP) was elevated in the anterior midgut, but not posterior midgut (Figure 4E). Together, these results confirm previous observations that *A. pomorum* increases lysine acetylation in *Tk+* EEs, suggesting possible epigenetic regulation or priming of *Tk* regulation, but correspondence between lysine acetylation and *Tk* reporter expression appears specific to certain regions of the midgut.

**Figure 4.**
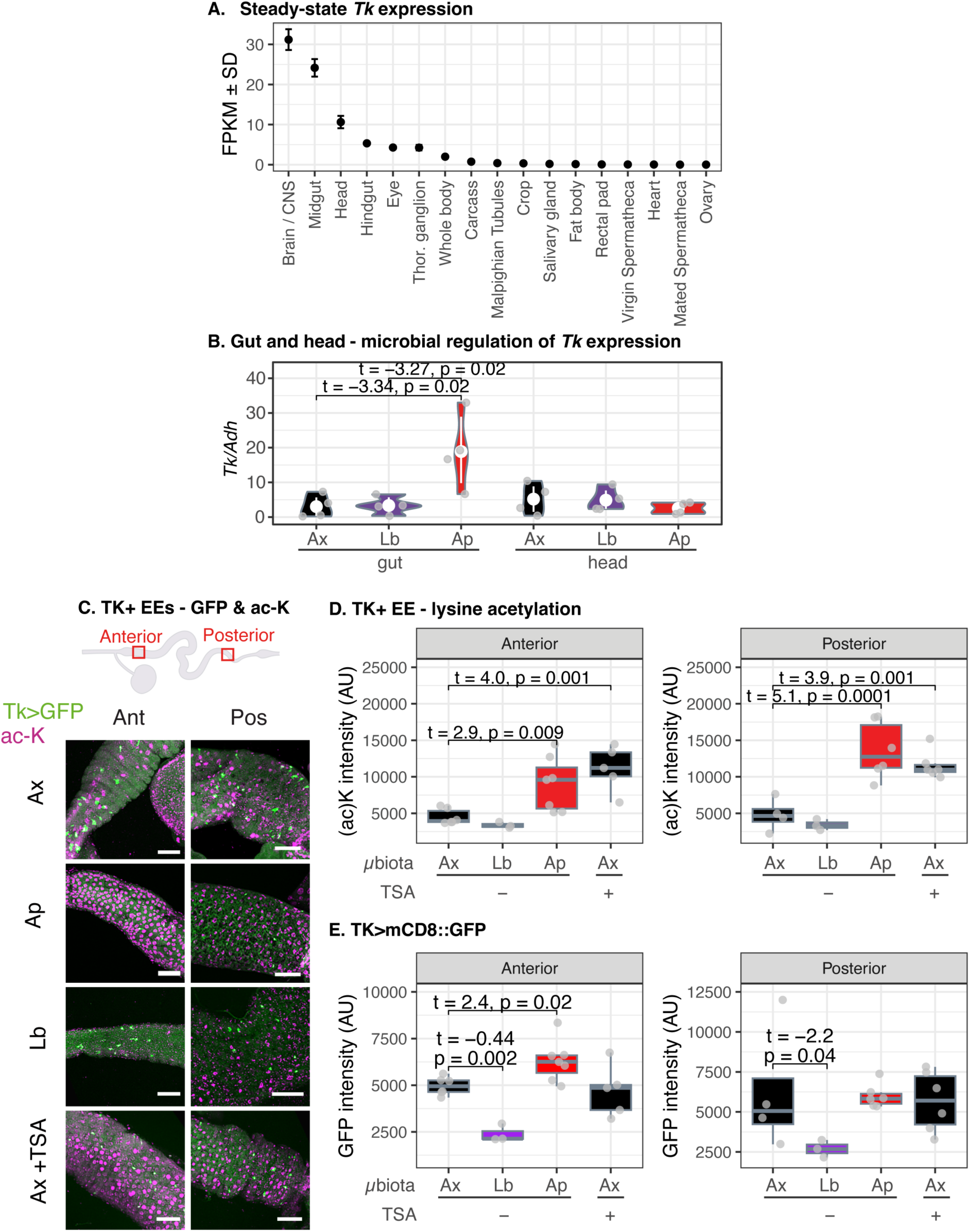
*A. pomorum* modulates intestinal *Tk* expression. **A.** *Tk* expression is enriched in nervous system and midgut tissues (data from FlyAtlas2). **B.** RT-qPCR of *Tk* in dissected midguts and heads of axenic and gnotobiotic flies indicates that expression is increased specifically in the midgut by *A. pomorum,* but not in the head; while *L. brevis* affects expression in neither tissue. White point and error bars show mean±SD. Statistics on brackets are from *post-hoc* EMM analyses of linear models, when significant differences were detected. **C.** Quantifying lysine acetylation and *Tk* promoter activity in *Tk*+ cells. Cartoon shows regions quantified in *Tk-Gal4/UAS-mCD8::GFP* flies with acetyl-lysine staining. Scale bars=50µm. **D.** Lysine acetylation is increased specifically by *A. pomorum* in *Tk+* enteroendocrine cells. Each point represents average fluorescence intensity for acetyl-lysine staining of ≥3 cells (marked by *Tk>GFP*) per gut, in anterior and posterior regions. The HDAC inhibitor Trichostatin-A was fed to axenic flies to confirm that increased histone acetylation is detectable by total acetyl-lysine staining. Statistics from one-way ANOVA, showing comparisons relative to axenic control condition. **E.** *A. pomorum* increases *Tk* promoter activity (*Tk-T2A-Gal4>UAS-GFP*) in anterior but not posterior midgut. Each point represents average GFP intensity of ≥3 cells per gut, in anterior and posterior regions. Statistics from one-way ANOVA, showing comparisons relative to axenic control condition.

Having shown that *Tk* expression was activated specifically by *A. pomorum* in the midgut, but not by *L. brevis*, we focussed specifically on *A. pomorum* in subsequent experiments.

### Intestine-directed *Tk* knockdown abrogates lifespan impact of *A. pomorum*

Next we asked whether *Tk* knockdown in the gut blocked the shortened lifespan of *Ap-*flies, using the *Gal4-UAS* system. In the midgut, EEs are marked by *voila-Gal4*, but this construct is also expressed in some neurons (Lin et al., 2022; Scopelliti et al., 2014), posing a challenge for fully parsing effects of neurons versus midgut EEs. We used a recently-developed intersectional strategy, in which *ChAT-Gal80* represses neuronal *voila-Gal4* (Medina et al., 2024), directing Gal4 to the midgut while sparing CNS. After confirming that this system was effective in our lab’s genetic background (Figure S8), we tested whether intestine-directed *Tk^RNAi^* abrogated impact of *A. pomorum* on lifespan, including every possible combination of *Voila-Gal4*, *ChAT-Gal80*, and *UAS-Tk^RNAi^* in a fully-factorial design (Figure 5A-B). Without *Tk^RNAi^*, lifespan was reduced in *Ap*-flies relative to axenic controls. However, expressing *Tk^RNAi^* in all *voila*+ cells (without *ChAT-Gal80*) blocked this effect of *A. pomorum* on lifespan. *ChAT-Gal80* restored a small effect of microbiota, but the magnitude of effect (Z-score) was diminutive, only ∼27-40% of that observed in controls. A second CoxPH analysis of the same data confirmed an overall 4-way interaction of *A. pomorum* and the three transgenes, confirming that the impact of *A. pomorum* was contingent on activation of *Tk^RNAi^,* and the tissues in which it was activated (p=0.0005). These analyses indicated that *Tk* in *Voila+* cells was obligately required for *A. pomorum* to modulate lifespan, and that the majority of this effect was explained by the intestine.

**Figure 5.**
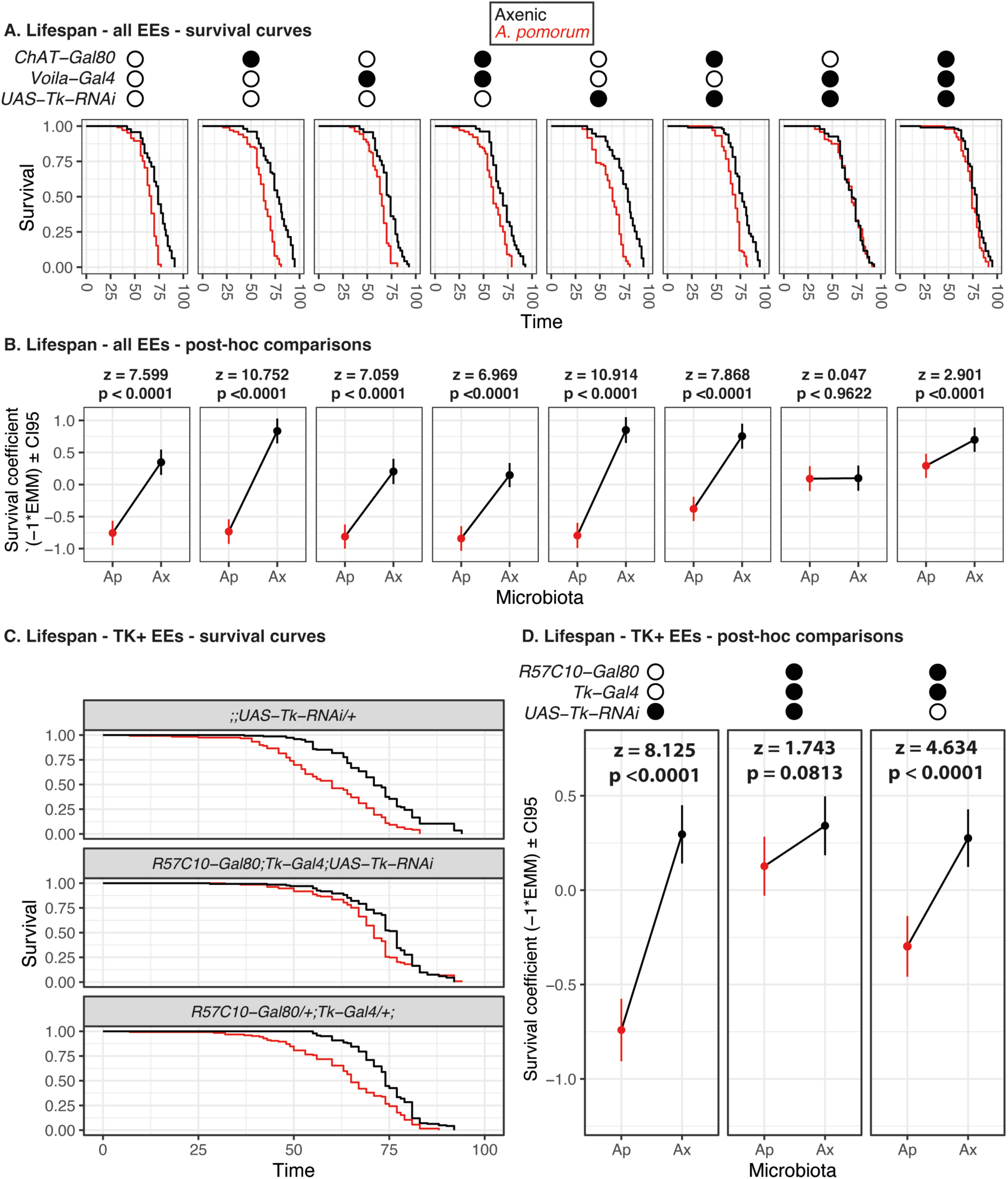
Gut-directed *Tk^RNAi^* abrogates lifespan shortening by *A. pomorum*. **A.** Kaplan-Meier survival plots of the indicated genotypes in presence/absence of *A. pomorum*, with knockdown activated by *Voila-Gal4,* in presence/absence of *ChAT-Gal80*. Statistics from *post-hoc* tests of CoxPH model. Genotypes indicated above panels. See Figure S8 for tissue-specificity of transgene combinations. All conditions n=105. **B.** *Post-hoc* analysis of survival model reveals attenuated lifespan response to *A. pomorum* with midgut-directed *Tk^RNAi^* (Genotypes as in A). Text indicates differences between *A. pomorum* and axenic flies in the given genotype. CoxPH analyses detected significant interactions both for microbiota*genotype (p<2.2e-16) and for Microbiota*:Voila-Gal4:ChAT-Gal80:UAS-TkRNAi* (p=0.0005). **C.** Kaplan-Meier survival plots of the indicated genotypes in presence/absence of *A. pomorum*, with knockdown activated by *Tk-T2A-Gal4,* and neuronal activity suppressed by *R57C10-Gal80*. **D.** *Post-hoc* analysis of survival model reveals attenuated lifespan response to *A. pomorum* with *Tk^RNAi^* directed to midgut by *Tk-T2A -Gal4* and *R57C10-Gal80*. Statistics from *post-hoc* tests of CoxPH model, indicating differences between *Ap*-flies and axenic flies in the given genotype. CoxPH detected overall genotype-by- *A. pomorum* interaction (p=2.46e-5).

For further validation we conducted a second lifespan experiment, using independent genetic tools (Rewitz et al., 2024) to direct *Tk^RNAi^* specifically to *Tk*+ EEs, with *Gal4* expressed from the *Tk* locus (*Tk-T2A-Gal4*), recombined with a transgene expressing *Gal80* under control of a fragment of *nSyb* promoter (*R57C10-Gal80*), which is expected to lead to pan-neuronal *Gal80* expression as another means to attenuate neuronal Gal4 activity (Kubrak et al., 2022; Malita et al., 2022; Rewitz et al., 2024). We again detected an *A. pomorum*-by-genotype effect (Figure 5C-D), with the lifespan-shortening effect of *A. pomorum* attenuated by *Tk* knockdown, and post-hoc tests showing only a marginal effect of *A. pomorum* on lifespan upon *Tk^RNAi^* induction (p=0.08). Independent *Gal4* and *Gal80* transgenes therefore corroborated the finding that intestinal *Tk* is required for the majority of *A. pomorum*’s lifespan-shortening effect.

### Role for neuronal *TkR99D^RNAi^* in host ageing effects of *A. pomorum*

We asked which receptors might mediate the *Tk-*dependent response to *A. pomorum*, and from which tissues. The fly genome expresses two cognate Tk receptors (TkRs) - *TkR86C* and *TkR99D -* which have different affinities for the six peptides encoded by *Tk* (Nässel et al., 2019), and distinct physiological roles. Plotting steady-state *TkR86C* and *TkR99D* expression, again using FlyAtlas2 (CR fly) data (Leader et al., 2018), revealed that both TkRs were most strongly expressed in CNS (Figure 6A). We expected that knocking down either one or both TkRs should extend lifespan in *Ap*-flies, mimicking effects of *Tk* knockdown.

**Figure 6.**
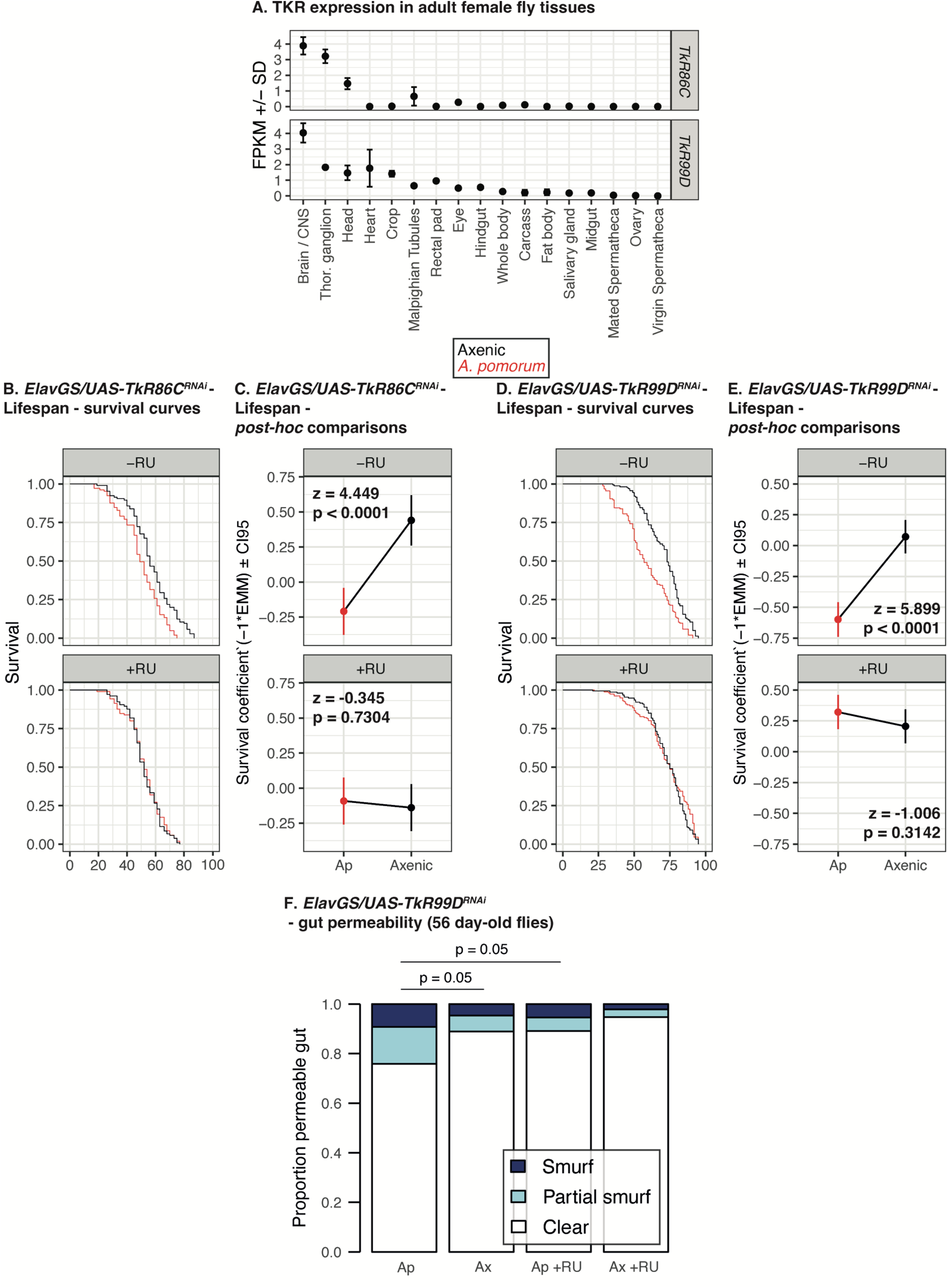
Pan-neuronal *TkR99D^RNAi^* abrogates lifespan impact of *A. pomorum*. **A.** FlyAtlas data showing expression of the two cognate *Drosophila* Tk receptors, *TkR86C* and *TkR99D*, among adult female tissues. **B.** Kaplan-Meier survival plots of *Elav-GS; UAS-TkR86C^RNAi^* flies in presence/absence of *A. pomorum*, with knockdown activated by RU486 feeding (bottom facet). CoxPH detected significant microbiota*RU interaction (p=0.0005). All conditions n=105. **C.** Post-hoc analysis of data from B showing that *TkR86C^RNAi^* induction locks axenic flies into a shortened lifespan. Text indicates differences between *A. pomorum* and axenic flies in each given RU condition. **D.** Kaplan-Meier survival plots of *Elav-GeneSwitch; UAS-TkR99D^RNAi^* flies in presence/absence of *A. pomorum*, with knockdown activated by RU486 feeding (bottom facet). CoxPH detected significant microbiota*RU interaction (p=1.362e-6). Ap RU- n=163, Ap RU+ n=161, Axenic RU- n=165, Axenic RU+ n=164 (pool of two replicate experiments). **E.** Post-hoc analysis of data from B showing that *TkR99D^RNAi^* induction blocks shortening of lifespan by *A. pomorum*. Text indicates differences between *A. pomorum* and axenic flies in each given RU condition. **F.** Neuronal *TkR99D^RNAi^* blocks onset of gut permeability (“smurf” phenotype) in aged flies. Bars show relative frequencies of flies with blue dye appearing to permeate through all tissues (“full smurf”) and through abdomen/thorax (“partial smurf”). χ^2^=16.126, DF=6, p=0.013. P-values on figure from chi-square tests of each given pair of columns, shown when differences were observed (otherwise omitted).

We tested the lifespan impacts of adult-onset TkR manipulation. RNAi constructs were developmentally lethal when crossed to ubiquitous drivers (*DaGS* and *ActGS*), even in the absence of inducer (RU), so instead we established lifespan experiments using the pan-neuronal driver, *Elav-GS*. Lifespan was shorter in *Ap-*flies relative to controls, and RNAi against each TkR interacted with the lifespan effect of *A. pomorum*, but in notably distinct ways. *TkR86C^RNAi^* had no effect in *Ap*-flies, but axenics were no longer long-lived (Figure 6B-C): this reveals an interaction with *A. pomorum* but does not recapitulate that of *Tk^RNAi^,* suggesting that TkR86C likely does not signal downstream of the *A. pomorum*-*Tk* relay. By contrast, *TkR99D^RNAi^* extended lifespan in *Ap-*flies, equivalent to axenics (Figure 6D-E). The most parsimonious explanation of these results is an *Acetobacter*-Tk-TkR99D relay, which shortens lifespan.

In fly midgut, TkR99D reportedly plays a role in mobilising lipid stores in enterocytes of the fly midgut (Song et al., 2014), prompting us to ask whether it may play an additional role in lifespan from these cells. However, expressing *TkR99D^RNAi^* with the enterocyte driver *Mex-GS* did not extend *Ap*-flies lifespan, suggesting that the *A. pomorum-*Tk relay signals gut-nonautonomously (Figure S9).

Finally, we asked whether lifespan extension in axenics and upon knockdown of *TkR99D* was accompanied by improved health in older flies (i.e. healthspan). Fly microbiota compromise healthspan by promoting gut barrier dysfunction in aged animals, characterised by permeability which shortly precedes death (Clark et al., 2015; Regan et al., 2016; Rera et al., 2012; Zane et al., 2023). We therefore tested whether *A. pomorum* promoted gut barrier dysfunction, and whether this could be rescued by *TkR99D^RNAi^* induction. We aged axenic and *Ap*-flies, with *TkR99D^RNAi^* from day 3 of adulthood onwards, before feeding on a blue dye to report gut permeability after 56 days (Figure 6F). *Ap-*flies indeed exhibited a higher rate of gut permeability than axenics, but *TkR99D^RNAi^* rescued the pathology to a level that was not significantly different from axenics. Thus, *TkR99D^RNAi^* can ameliorate deleterious impacts of *A. pomorum* on late-life health, suggesting that the putative *A. pomorum*-*Tk-TkR99D* relay limits late-life health.

### Lifespan and metabolic impacts of *A. pomorum* are not explained by *TkR99D* in insulin-producing cells

What happens downstream of the putative *A. pomorum-*Tk-TkR99D relay? Reduced insulin signalling promotes healthy ageing across species, with lifespan and other benefits reported in flies, nematodes and rodents, and genetic associations indicating potential conservation in humans (Clancy et al., 2001; Kenyon et al., 1993; Selman et al., 2008; Slagboom et al., 2017). *A. pomorum* is reported to activate insulin signalling (S. Shin et al., 2011), while *TkR99D* expression is reported in insulin-producing cells (IPCs) in the fly brain (Birse et al., 2011), ablation of which extends lifespan (Alic et al., 2014; Broughton et al., 2005). We therefore hypothesised that Tk-TkR99D signalling impacts ageing by serving as a relay between the gut microbiota and IPCs (Figure 7A).

**Figure 7.**
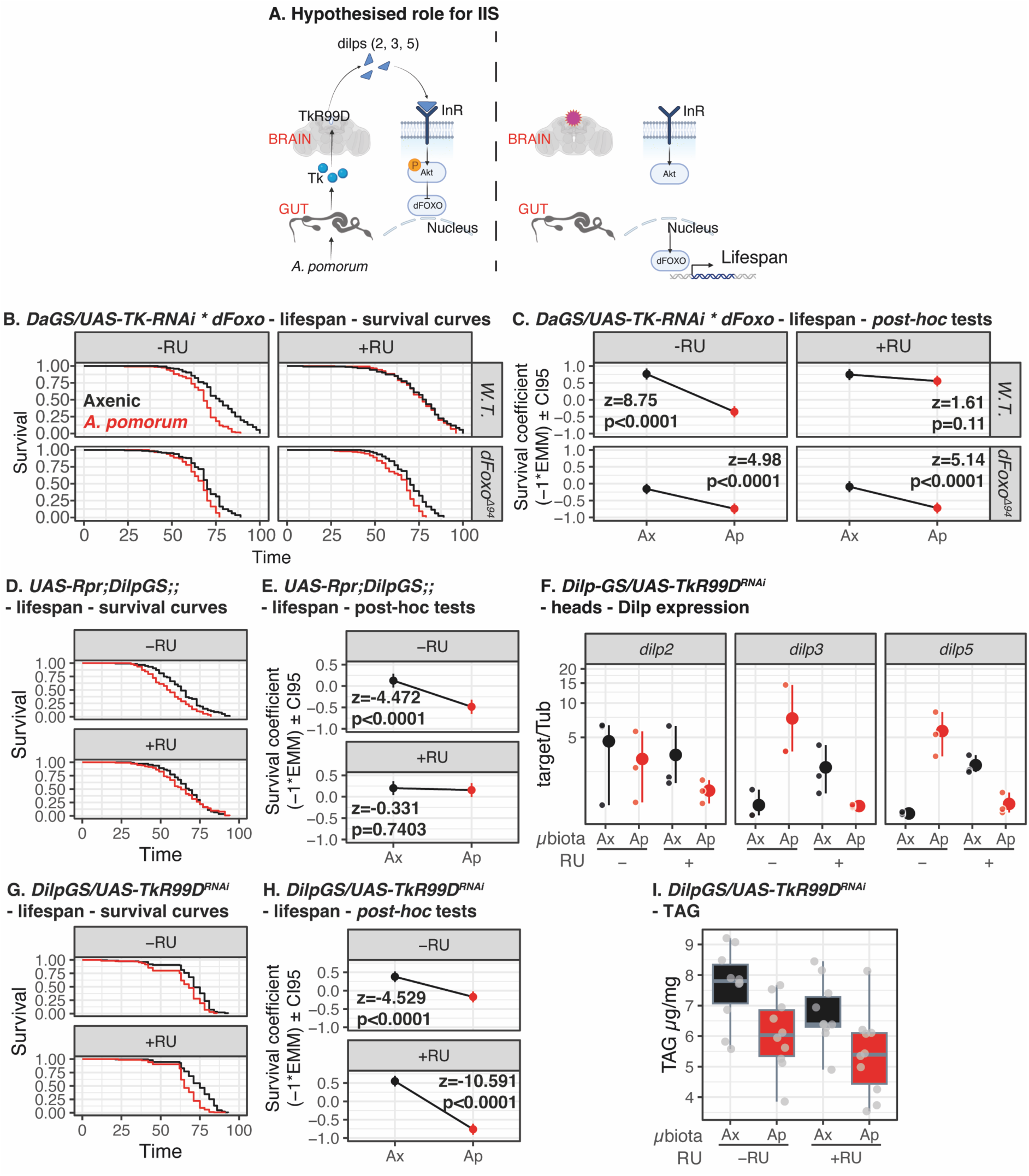
Insulin signalling is involved in lifespan response to *A. pomorum* and *TkR99D* knockdown, but not obligately required. **A.** Hypothetical model of role for insulin signalling. In presence of microbiota (left), Tk peptides are released from gut and contact TkR99D in IPCs/brain. IPCs are released into circulation, activating insulin signalling in peripheral tissues and leading to nuclear exclusion of dFOXO. In absence of microbiota (right), extracellular signalling is diminished, reducing downstream insulin signalling and consequent activation of lifespan-extending gene expression program by dFOXO. **B.** dFOXO connects lifespan responses to *A. pomorum* and ubiquitous *Tk^RNAi^*. Facets show Kaplan-Meier plots in axenic and *A. pomorum* conditions, with/without RNAi induction (RU), and with/without *dFoxO^Δ94^* null mutation. Overall microbiota*RU*Foxo interaction p=5.384e-05 (CoxPH). All conditions n=150 except W.T. *A. pomorum* -RU (n=135) and W.T. axenic +RU (n=148). **C.** Post-hoc comparisons (EMmeans), showing impact of *A. pomorum* under specified conditions of *TkRNAi* induction and *dFoxO* deletion. *A. pomorum* shortens wild-type lifespan, but this effect is diminished by *dFoxO^Δ94^. Tk^RNAi^* induction blocks lifespan shortening by *A. pomorum* in wild-type background, but not in *dFoxO^Δ94^* background; altogether suggesting that *dFoxO* is required for *A. pomorum* to influence longevity via Tk modulation. **D-E.** Insulin-producing cells (IPCs) are required for *A. pomorum* to influence lifespan (CoxPH p=0.003). All conditions n=120. Post-hoc tests (**E**) confirm that lifespan effect of *A. pomorum* (-RU) is absent following IPC ablation (*Rpr* expression, +RU). **F.** *A. pomorum* increases expression of *Dilp3* and *Dilp5,* but not *Dilp2*, in a *TkR99D -*dependent manner. Panels show expression (RT-qPCR) in heads of axenic and *Ap*-flies ±*TkR99D^RNAi^* in IPCs (*Dilp2-GS/UAS-TkR99D^RNAi^*). by Microbiota*RU interactions were tested for with linear models: *Dilp2* F_1,24_=0.424, p=0.5, *Dilp3* F_1,24_=23.404, p=0.0001, *Dilp5* F_1,24_=29.261, p<0.0001. **G-H.** *TkR99D^RNAi^* in IPCs worsens lifespan impact of *A. pomorum*. Ap -RU n=160, Ap +RU n=162, Ax -RU n=160, Ax +RU n=163. (**I**) Post-hoc tests indicate accentuated shortening of lifespan by *A. pomorum* following *TkR99D^RNAi^* induction (+RU). **I.** *TkR99D^RNAi^* in IPCs does not modulate metabolic impact of *A. pomorum* on fly TAG. The two experimental factors each had significant effects on TAG (ANOVA, *A. pomorum* F=14.95, Df=1,37, p=0.0004; RU F=4.32, Df=1,37, p=0.04), but no interaction was detected.

We tested whether insulin signalling was required for lifespan to respond to *A. pomorum*. The forkhead box-O transcription factor dFOXO, encoded by the *dFoxO* locus, is obligately required for longevity upon reduced insulin signalling (Slack et al., 2015, 2011). dFOXO is retained in cytoplasm of Ap-flies, consistent with activated insulin signalling (S. Shin et al., 2011). We recombined a dFOXO null mutation (*dFoxO^Δ94^*) with transgenes to knock down *Tk* ubiquitously, performed lifespan assays, and used Cox proportional hazards analysis to test for an *A. pomorum*Tk*dFoxO* interaction (Figure 7B-C). We observed a significant three-way interaction (CoxPH p=5.384e-05) – indicating that *dFoxO^Δ94^* modulated the microbiota-*Tk* interaction – and then used post-hoc tests to dissect the co-dependence of the three experimental variables (Table 1). In controls without *Tk* knockdown, *A. pomorum* shortened lifespan relative to axenics in wild-type flies, but *dFoxO^Δ94^* flies exhibited a diminished but still-significant shortening of lifespan by *A. pomorum*. Without *dFoxO^Δ94^, Tk* knockdown blocked lifespan response by rendering *Ap*-flies long-lived. In the *dFoxO^Δ94^* background*, Tk* knockdown did not extend lifespan, in either *Ap-*flies or axenics. However, unexpectedly, *Tk^RNAi^* ceased to block the effect of *A. pomorum* in the *dFoxO^Δ94^* background. Thus, in summary, (A) *dFoxO* is partially required for lifespan effect of *A. pomorum*, (B) *dFoxO* is required for lifespan extension upon *Tk* knockdown, and (C) *dFoxO* is required for *A. pomorum* and *Tk* knockdown to have an interactive effect on lifespan. We interpret that *dFoxO* potentiates the putative *A. pomorum-*Tk-TkR99D relay.

**Table 1.** *dFoxO* is partially required for microbiota\**Tk* interaction for lifespan: post-hoc pairwise comparisons (EMMeans)

| contrast | estimate | SE | z.ratio | p.value |
| --- | --- | --- | --- | --- |
| (Ax WT RU-) - (Ap WT RU-) | -1.12 | 0.13 | -8.76 | 6.38E-14 |
| (Ax WT RU-) - (Ax dFoxOΔ RU-) | -0.92 | 0.12 | -7.51 | 1.75E-12 |
| (Ax WT RU-) - (Ap dFoxOΔ RU-) | -1.51 | 0.13 | -11.87 | 0 |
| (Ax WT RU-) - (Ax WT RU+) | -0.02 | 0.12 | -0.15 | 0.9999999 |
| (Ax WT RU-) - (Ap WT RU+) | -0.21 | 0.12 | -1.77 | 0.63954118 |
| (Ax WT RU-) - (Ax dFoxOΔ RU+) | -0.85 | 0.13 | -6.79 | 3.21E-10 |
| (Ax WT RU-) - (Ap dFoxOΔ RU+) | -1.48 | 0.13 | -11.65 | 0 |
| (Ap WT RU-) - (Ax dFoxOΔ RU-) | 0.2 | 0.12 | 1.66 | 0.71153814 |
| (Ap WT RU-) - (Ap dFoxOΔ RU-) | -0.39 | 0.12 | -3.16 | 0.03369588 |
| (Ap WT RU-) - (Ax WT RU+) | 1.1 | 0.13 | 8.6 | 7.16E-14 |
| (Ap WT RU-) - (Ap WT RU+) | 0.91 | 0.13 | 7.27 | 1.00E-11 |
| (Ap WT RU-) - (Ax dFoxOΔ RU+) | 0.27 | 0.13 | 2.13 | 0.39332695 |
| (Ap WT RU-) - (Ap dFoxOΔ RU+) | -0.36 | 0.12 | -2.96 | 0.06098085 |
| (Ax dFoxOΔ RU-) - (Ap dFoxOΔ RU-) | -0.59 | 0.12 | -4.99 | 1.70E-05 |
| (Ax dFoxOΔ RU-) - (Ax WT RU+) | 0.9 | 0.12 | 7.34 | 5.87E-12 |
| (Ax dFoxOΔ RU-) - (Ap WT RU+) | 0.71 | 0.12 | 5.93 | 8.56E-08 |
| (Ax dFoxOΔ RU-) - (Ax dFoxOΔ RU+) | 0.06 | 0.12 | 0.54 | 0.99943704 |
| (Ax dFoxOΔ RU-) - (Ap dFoxOΔ RU+) | -0.57 | 0.12 | -4.77 | 5.07E-05 |
| (Ap dFoxOΔ RU-) - (Ax WT RU+) | 1.49 | 0.13 | 11.7 | 0 |
| (Ap dFoxOΔ RU-) - (Ap WT RU+) | 1.3 | 0.12 | 10.45 | 7.78E-14 |
| (Ap dFoxOΔ RU-) - (Ax dFoxOΔ RU+) | 0.65 | 0.12 | 5.35 | 2.43E-06 |
| (Ap dFoxOΔ RU-) - (Ap dFoxOΔ RU+) | 0.02 | 0.12 | 0.19 | 0.99999961 |
| (Ax WT RU+) - (Ap WT RU+) | -0.19 | 0.12 | -1.61 | 0.74370666 |
| (Ax WT RU+) - (Ax dFoxOΔ RU+) | -0.84 | 0.13 | -6.63 | 9.28E-10 |
| (Ax WT RU+) - (Ap dFoxOΔ RU+) | -1.47 | 0.13 | -11.49 | 0 |
| (Ap WT RU+) - (Ax dFoxOΔ RU+) | -0.64 | 0.12 | -5.23 | 4.63E-06 |
| (Ap WT RU+) - (Ap dFoxOΔ RU+) | -1.27 | 0.12 | -10.23 | 6.75E-14 |
| (Ax dFoxOΔ RU+) - (Ap dFoxOΔ RU+) | -0.63 | 0.12 | -5.14 | 7.52E-06 |

To assess directly the role of tissues reported to express *TkR99D* (Birse et al., 2011), we turned to the IPCs. We first tested whether ablating these cells by expressing the pro-apoptotic factor *Rpr* would block lifespan response to *A. pomorum*, reasoning that a role for these cells in response to the bacteria would be shown by a lack of response following ablation. Lifespan was again shortened in *Ap-*flies relative to axenic controls. However this effect was blocked by adult-onset *Rpr* expression in IPCs (using the *Dilp2-GS* driver) (Figure 7D-E), and we detected an overall microbiota*RU interaction (CoxPH p=0.003). This indicated a role for IPCs in lifespan shortening by *A. pomorum*. To test whether TkR99D activation could account for this effect we expressed *TkR99D^RNAi^* in IPCs, and assayed expression of the three insulin-like peptides expressed therein (*Ilp2, Ilp3, Ilp5*) by RT-qPCR of heads (Figure 7F, Table 2). *Ilp2* exhibited no response to *A. pomorum* or *TkR99D^RNAi^*.(linear model, microbiota:RU interaction F_1,24_=0.424, p=0.5) However, for *Ilp3* and *Ilp5*, *A. pomorum* promoted expression, and this effect was blocked by *TkR99D^RNAi^* induction (linear models, microbiota:RU interactions *Ilp3* F_1,24_=23.404, p=0.0001, *Ilp5* F_1,24_=29.261, p<0.0001.) Specifically, *Ilp3* and *Ilp5* expression levels were increased in *Ap-*flies, but these effects were absent upon *TkR99D^RNAi^* induction (Table 2).

**Table 2.**
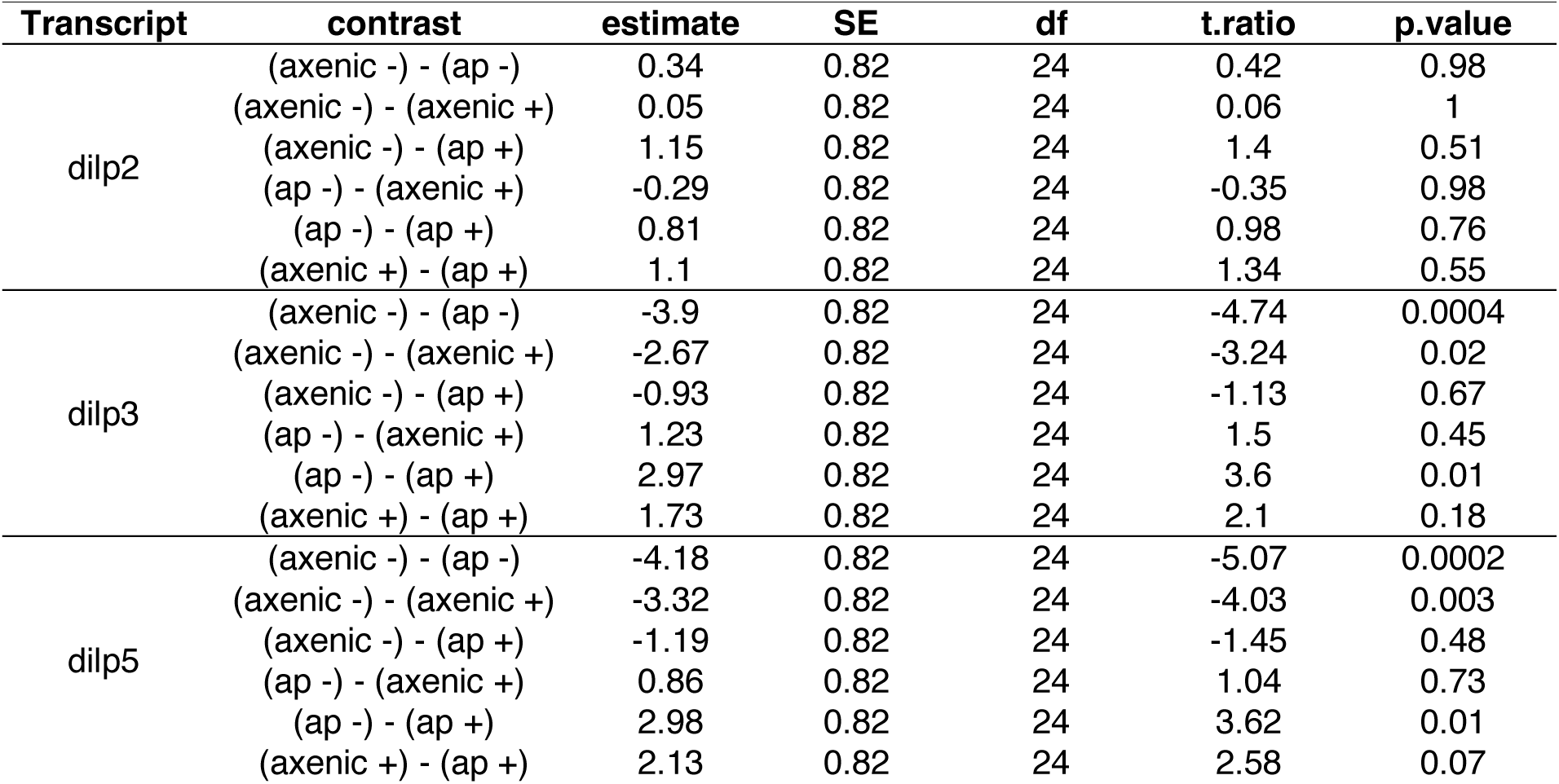
*TkR99D^RNAi^* in insulin-producing cells blocks increased expression of *dilp3* and *dilp5*, but not *dilp2*, by *A. pomorum*: *post-hoc* tests.

Finally, we tested whether *TkR99D^RNAi^* in IPCs blocked lifespan shortening by *A. pomorum* (Figures 7G-H). As expected, without *TkR99D^RNAi^, Ap*-flies were shorter-lived than axenics. *TkR99D^RNAi^* induction in IPCs did not affect axenic lifespan but, surprisingly, the lifespan-shortening effect of *A. pomorum* was exaggerated by *TkR99D^RNAi^* induction (Figure 7H). Consequently, *A. pomorum’s* effect on lifespan seemingly cannot be explained by TkR99D activation in IPCs. We also tested whether *TkR99D^RNAi^* in IPCs altered effects of *A. pomorum* on TAG levels in young flies (Figure 7I): no interaction was apparent (though we detected independent effects of microbiota (ANOVA F=14.95, Df=1,37, p=0.0004) and RNAi induction (F=4.32, Df=1,37, p=0.04)), suggesting that interactive effects of *A. pomorum* and Tk peptides are not mediated by *TkR99D* in IPCs.

Altogether, these data outline involvement of insulin signalling in responses to *A. pomorum* and *Tk*, because IPC ablation blocked lifespan shortening by *A. pomorum* (Figure 7D-E), and because *dFoxO* was required for the interaction of *A. pomorum* and *Tk^RNAi^*. However, the failure of *TkR99D^RNAi^* in IPCs to block *A. pomorum’s* effects on lifespan or TAG suggests that insulin signalling does not act downstream of the putative *A. pomorum*/Tk/TkR signalling relay. These data may suggest priming, with insulin signalling required to potentiate responses to *A. pomorum*/Tk/TkR.

### No evidence of role for *TkR99D* in Akh-producing cells in lifespan and metabolic response to *A. pomorum*

In insects, adipokinetic hormone (Akh) is released from *corpora cardiaca* (cc) to promote lipid mobilisation in response to heightened energy demand, analogous to glucagon in mammals (Gäde and Marco, 2022; Musselman et al., 2018; Toprak et al., 2020). Having observed *Tk*-dependent effects of microbiota on TAG storage, and following recent evidence that *TkR99D* in Akh-producing cells limits lifespan (Ahrentløv et al., 2025), we tested whether it may also mediate lifespan impact of *A. pomorum*.

First, to test whether Akh peptide levels in CC were *A. pomorum*-dependent, we quantified Akh with antibody staining. Akh staining levels were significantly higher in CC dissected from *Ap-*flies than axenics (Figure 8A), suggesting microbial regulation of Akh signalling.

**Figure 8.**
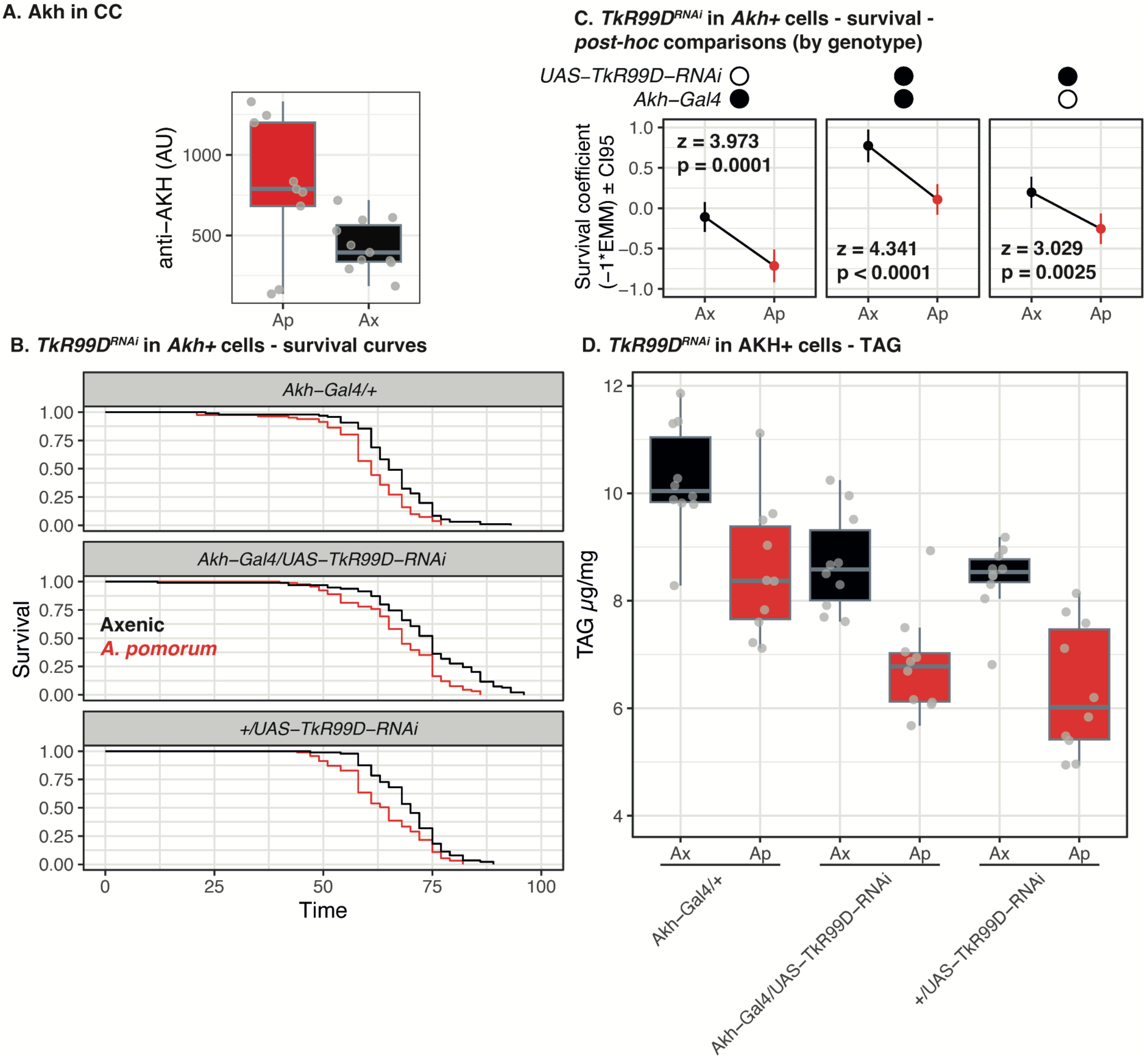
*A. pomorum* modulation of host lifespan and TAG is independent of *TkR99D* in Akh-producing cells. **A.** Antibody staining of Akh in dissected Corpora Cardiaca (CC) is increased in *Ap-*flies (Ap) relative to axenics (Ax). (t-test: t = −2.36, p-value = 0.04). **B.** Microbial shortening of lifespan is independent of *TkR99D* signalling in Akh+ cells. Facets show Kaplan-Meier survival curves for axenic and *Ap-*flies when *TkR99D* is knocked down (middle facet) and in each control genotype. No significant interaction of genotype and *A. pomorum* was detected (CoxPH p=0.58). *Akh-Gal4/+ A. pomorum* n=103, *Akh-Gal4/+* axenic n=108, *Akh-Gal4/UAS-TkR99D-RNAi A. pomorum* n=105, *Akh-Gal4/UAS-TkR99D-RNAi* axenic n=107, *+/UAS-TkR99D-RNAi A. pomorum* n=103, *+/UAS-TkR99D-RNAi* axenic n=101. **C.** Post-hoc comparisons (EMmeans) confirm equivalent lifespan shortening by *A. pomorum*, independent of *TkR99D^RNAi^*. Combination of transgenes indicated above facets. **D.** *A. pomorum* modulation of host TAG is independent of *TkR99D^RNAi^* in Akh+ cells. Boxplots show TAG levels are reduced independent of *TkR99D^RNAi^* induction. No significant interaction of genotype and *A. pomorum* detected.

We then tested directly whether TkR99D was required in Akh-producing cells for *A. pomorum* to shorten lifespan. *A. pomorum* shortened lifespan across the all different *Akh* conditions (Figure 8B-C). Expressing *TkR99D^RNAi^* under control of *Akh-Gal4* did extend lifespan in both axenics and *Ap-*flies, corroborating a recent report of the CC’s role in longevity (Ahrentløv et al., 2025) (noting background effects, despite backcrossing immediately before experimentation) (Figure 8B-C). However, these data indicate that lifespan shortening by *A. pomorum* is independent of TkR99D in CC. We also tested whether RNAi against Akh receptor (*AkhR*) in adipose (fat body) altered response to *A. pomorum*, driving expression with *S106-GS*, but again no interaction was apparent (Figure S10). Finally, we tested whether *TkR99D* in Akh-producing cells interacted with effects of *A. pomorum* on TAG storage. *TkR99D^RNAi^* induction did not block response to *A. pomorum*, with *Ap-*flies storing less lipid than axenics independent of fly genotype (Figure 8D), indicating that *A. pomorum*’s influence on TAG does not depend on TkR99D in CC. Combined with our investigation of insulin signalling, our studies of Akh suggest that the impact of *Acetobacter* on ageing is mediated by non-canonical mechanisms that depend on *Tk* and *TkR99D*.

## Discussion

Microbiota impact ageing across animals (Cabreiro et al., 2013; Cui et al., 2019; Sannino et al., 2018; Seidel and Valenzano, 2018; Smith et al., 2017; Snyder et al., 1990; Tazume et al., 1991), and connections between gut and the nervous system are key to health throughout the lifecourse (Boehme et al., 2022; Dinan and Cryan, 2017; Qansuwa et al., 2024). Here we have connected microbial regulation of ageing with tissue-specific regulation of endocrine signalling genes in gut and neurons. Our *Drosophila* results show that (A) *A. pomorum* is sufficient to modulate *Tk* in the midgut, (B) knocking down the peptide in midgut blocks the lifespan response to the bacteria, and (C) neuronal knockdown of the receptor *TkR99D* recapitulates these effects. We are struck by the specificity of this circuit, with knockdown of a single peptide-encoding locus in a specific tissue ablating the microbial impact on lifespan, despite the wide array of host functions that the microbiota affect. This specific requirement can likely be explained by the highly pleiotropic functions of Tachykinins (Nässel, 2025; Nässel et al., 2019), suggesting that *Tk* knockdown reprograms organismal physiology to a state in which lifespan can no longer be modulated by microbiota. Given the evolutionary conservation of tachykinins, it will be interesting in future work to ask whether this family of peptides can also modulate ageing in other species.

Our work indicates that the influence of the microbiota-*Tk/TkR99D* relay is partially independent of better-studied lifespan assurance mechanisms, i.e. insulin signalling, and phenotypes associated with ageing. The lifespan benefit of *Tk* knockdown was not accompanied by costs to fecundity or feeding rate in CR flies, but instead knockdown altered metabolic impact of microbiota. Furthermore, knocking down *TkR99D* in major signalling hubs associated with ageing and TAG metabolism (IPCs and Akh+ cells, respectively) did not block lifespan or TAG response to *A. pomorum*, suggesting that yet-uncharacterised neuroendocrine mechanisms are at play. We interpret the partial requirement for *dFoxO*, connecting *A. pomorum* to *Tk^RNAi^* modulation of lifespan, as suggesting that insulin potentiates downstream responsiveness, without necessarily being involved directly by *TkR99D* in IPCs.

Our work was performed entirely in females. We chose to study the sex that tends to exhibit more robust response to anti-ageing interventions (Regan and Partridge, 2013), because of the labour-intensive and technically challenging nature of performing lifespan experiments under lifelong axenic conditions. Having established this the *A. pomorum-Tk-TkR99D* connection, it will be interesting in future work to ask whether males also enjoy a lifespan benefit of diminishing the activity of this circuit.

*Tk* is one of a number of signalling circuits implicated downstream of the microbiota, for example JAK-STAT and JNK signalling (Buchon et al., 2009), IMD (Jugder et al., 2021; Watnick and Jugder, 2020), and the neuropeptide CNMamide (Kim et al., 2021). We do not propose that Tachykinins alone are sufficient to explain all physiological effects of the microbiota: rather we expect that the interplay of multiple signalling pathways collectively orchestrate the host organismal response to microbiota. Indeed our transcriptome (re)analysis (Figure 1B) suggested that a community of bacteria co-regulate expression of the neuropeptides *ITP, AstC* and *NPF* alongside *Tk.* We note that *Tk* and *NPF* are co-expressed in a number of the same EE populations (Guo et al., 2019), and related functions on nutrient-specific appetite, physiology and Akh signalling are reported (Ahrentløv et al., 2025; Malita et al., 2022), suggesting closely coupled functions for Tk and NPF which may also be relevant to integrating responses to microbiota. Further work should aim to identify the broader signalling networks that respond to the microbiota, and the extent to which the constituents and structure of those networks mirror what is known in other contexts (e.g. responses to diet or pathogenic infection).

A key element to better understanding how signalling networks at large respond to microbiota will be to better understand what properties of the microbiota the signals convey. Signalling is selected to convey information about environmental, nutritional, and physiological state: which of these parameters do the microbiota impact? Impacts on metabolism could be mediated by direct provision of nutrients (e.g. short-chain fatty acids, B-vitamins (Sannino et al., 2018; S. C. Shin et al., 2011; A. C.-N. Wong et al., 2014)), and/or competition with the host by removal of nutrients from diet (Huang and Douglas, 2015). Immunity may be modulated by bacterial titre, to prevent pathological overgrowth of otherwise mutualistic bacteria. Our results indicate that the interplay between bacteria and *Tk* knockdown is bacteria-specific, suggesting in turn that more complex mixes of bacteria that co-occur with the fly may elicit distinct signalling network topologies. Furthermore, the previous finding that fly transcriptional networks are structured by microbiota (Dobson et al., 2016) leads to a question of whether other host networks - such as endocrine signalling - may also be structured by microbiota, in which case signalling in the presence of one bacterium might elicit quite different physiological responses than in the presence of a distinct bacterium.

The two biggest mechanistic questions arising from this work are (A) from where in the nervous system does *TkR99D* inhibition extend lifespan? and (B) by what mechanism does inhibition extend lifespan? We tested two neuroendocrine tissues, the IPCs and the Akh+ cells, finding that *TkR99D^RNAi^* in either did not block shortened lifespan or TAG phenotypes in *Ap-*flies (though bacteria-independent effects of *TkR99D^RNAi^* in *Akh+* cells were apparent). These data suggest that we can reject a role for TkR99D in IPCs and *Akh+* cells, and instead that an elusive population of *TkR99D+* neurons is at play downstream of the *Acetobacter-Tk* relay. With respect to future mechanistic understanding, we suggest it will be important to note that pan-neuronal *TkR99D^RNAi^* also mitigated gut permeability (smurfing) in aged *Ap*-fies, because the smurf phenotype is underpinned by increased immune system activity (Clark et al., 2015; Clark and Walker, 2017), and activation of *Tk* cells by *Acetobacter* is dependent on immune (IMD) signalling (Jugder et al., 2021). Together these observations suggest close links between *Tk* signalling and immunity, outlining a possible role for *Acetobacter/Tk/TkR99D* in “inflammaging”.

The biggest comparative question arising from our work is whether our findings are relevant in other animals, potentially including mammals. Flies have a track record of revealing mechanisms of ageing that are conserved in other systems (Bjedov et al., 2010; Clancy et al., 2001; Harrison et al., 2009; Kenyon et al., 1993; Miller et al., 2014; Selman et al., 2008; Wilkinson et al., 2012; Zhang et al., 2024). Tachykinins are conserved from invertebrates to mammals, raising the question of whether their functions in ageing are also conserved. The next step would most likely be to assess whether Tachykinin knockdown has anti-ageing effects in rodents, and/or short-lived vertebrates like the African turquoise killifish. Such studies may pave the way for drug repurposing, as some small-molecule inhibitors of human TkRs have already been developed and are licenced in some jurisdictions for specific purposes. Such interest is timely since the therapeutic role of TkRs is receiving renewed attention for potential metabolic therapies (Sass et al., 2024).

In conclusion, our study establishes a mechanistic connection between the microbiota and ageing via gut/neuron signalling. Microbial shortening of host lifespan depends on *Tk/TkR* signalling, which can be explained by *Acetobacter* activating intestinal *Tk* and, downstream, neuronal *TkR99D*. The findings hint at effects which may be conserved in other systems, which may ultimately help to explain mechanistically why microbiota exert a conserved influence on animal health and ageing.

## Materials and methods

### Flies, bacteria, husbandry and culturing

Stocks used are detailed in Table 3. All stocks (except for recombinant *R57C10-Gal80;;Tk-T2A-Gal4,* gift from Kim Rewitz) were backcrossed into the Dahomey background (originally isolated from Benin in the 1970s), bearing the *w1118* mutation and confirmed *Wolbachia*-negative. Presence of *dFoxO^Δ94^* in recombinant flies was confirmed by PCR of mothers of single-fly crosses.

**Table 3.**
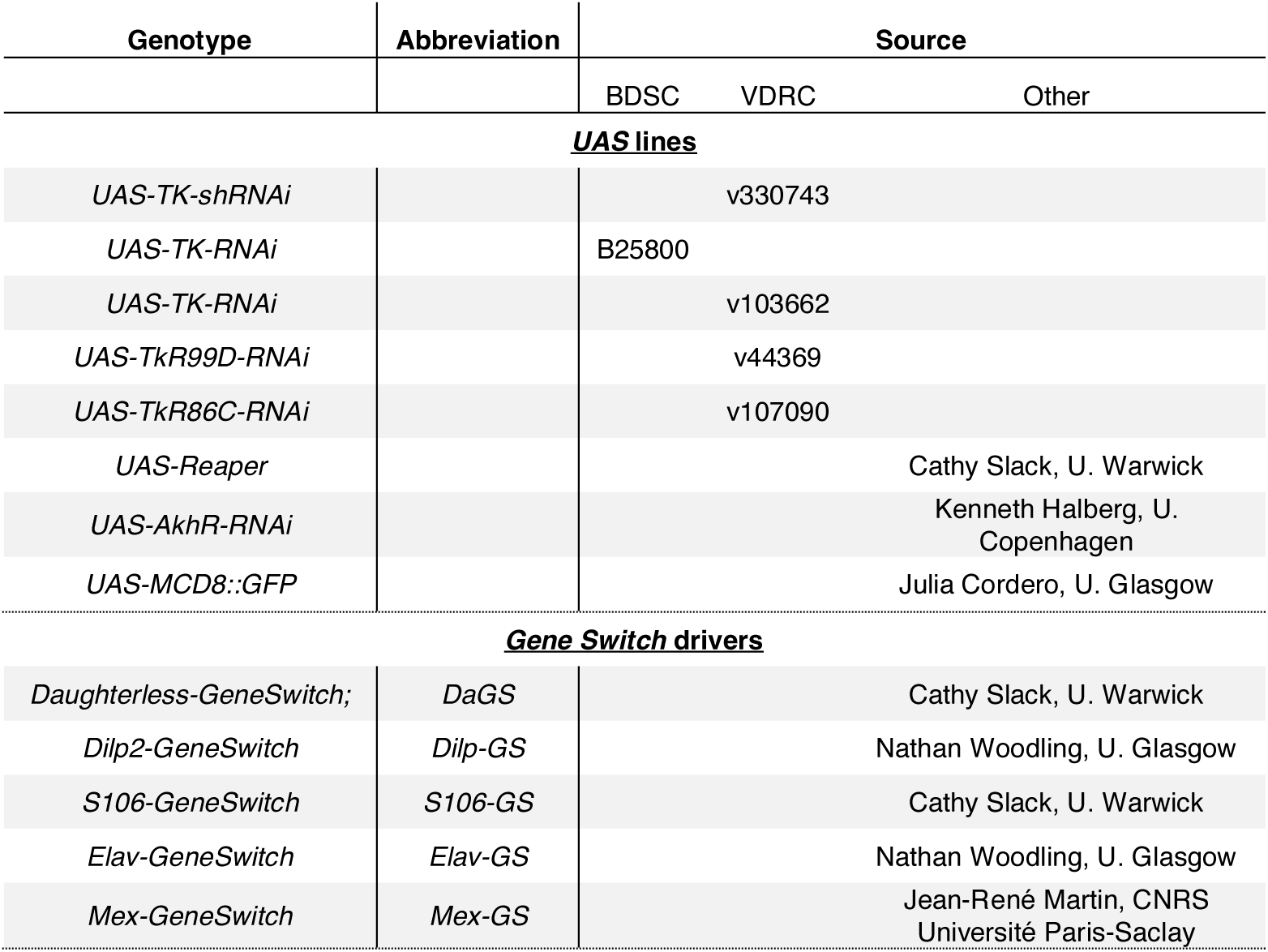
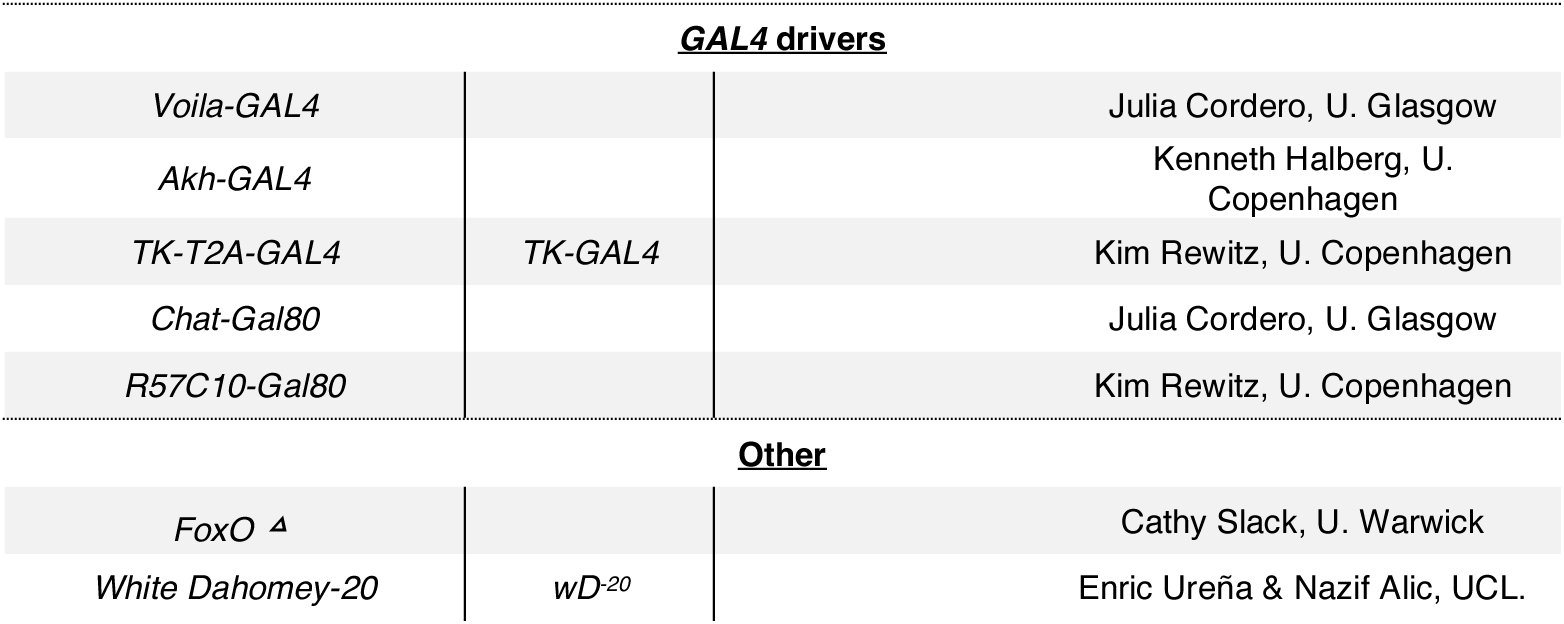
*Drosophila* stocks used in this study.

All flies were maintained at 25°C with a 12-12-hour light-dark cycle on SYA medium, comprising 5% sucrose (MP Biomedicals BP220), 10% yeast (MP Biomedicals, 903312, lot no. S7760), and 1.5% agar (Sigma-Aldrich A7002)(Bass et al., 2007; Sannino and Dobson, 2023). Preservatives (0.3% nipagin, 0.3% propionic acid) were incorporated into media for stock maintenance. Crosses were performed on juice agar in egg laying cages (per litre: 20g agar, 26g sucrose, 52g glucose, 7g yeast, 88.8ml organic clementine or grape juice). To generate axenic flies, eggs were collected and washed 2x in alternating 10% bleach for (3-5 min) and sterile ddH2O for (1 min), with an optional additional 1 min wash in EtOH, before being allowed to pellet by gravity in sterile PBS. 20 µl of egg pellet was aspirated with wide-mouthed pipette tips into T75 flasks containing 50mL SYA. Preservatives were included during fly development only in CR experiments. We made gnotobiotic embryos as as previously (Sannino and Dobson, 2023). M9+lactate media comprised 200ml 5x M9 salts, 2ml 1M MgSO_4_, 1ml 1M CaCl_2_, 25ml 20% (v/v) lactate, made up to 1l in ddH_2_O and autoclaved. YPD media comprised 10g yeast extract (VWR 84601.0500), 20g peptone (VWR 84620.0500), 20g dextrose (Fisher Life Sci G/0500/53). *Acetobacter pomorum* (DmCS004) was streaked and grown on M9+lactate plates (1.5% agar), before growing single colonies in liquid M9+lactate at 30°C with shaking (225 rpm) for 2 days. (DmCS004). *L. brevis* (DmCS003) was streaked on YPD (1.5% agar), then grown in liquid YPD for two days at 30C in static culture. Cultures were pelleted, washed, resuspended and diluted in sterile PBS to an OD_600_ of 0.2. 200 µL of bacterial suspensions were added to T75 flasks containing freshly-prepared axenic eggs.

Newly emerged flies transferred to fresh media using sterile technique in a laminar flow hood, (preservatives as per development) for the first three days of adulthood before gentle anaesthesia on CO_2_. Males were removed and females allocated to SYA containing preservatives in sterile glass *Drosophila* vials, using aseptic technique in a laminair flow hood with a sterile paint brush and sterile petri dish on ice. In experiments using RU486 (Mifepristone, Cayman Chemical, CAY10006317), 2ml l^− 1^ of 100mM RU486 stock solution in EtOH was incorporated into media at 200µM final concentration, and fed from day 3 adulthood onwards. 2ml EtOH was incorporated as vehicle control in RU-free controls.

For lifespan assays, mated females were allocated to sterile glassware in groups of 15, and transferred to fresh food every 2–3 days. At each transfer, deaths and censors were scored until all flies were dead. For axenic and gnotobiotic lifespans, fly tipping was performed in a sterile laminar flow hood with aseptic technique.

### Feeding and egg laying assays

Feeding rate was measured by quantifying average proboscis extension rate (PER) over 3-4 hours. Vials containing 5 mated females each were arranged on a benchtop in a 25°C room the night before behavioural observations to allow acclimation. Experimenters were blinded to vial ID and experimental condition. The following morning, PER was recorded by iteratively counting the instantaneous number of flies feeding per vial for 3-4 hours. The average of these measurements per each given vial was taken as that vial’s feeding rate. At the end of the experiment, flies were disposed, and vials frozen. The next day eggs were counted from the same frozen vials.

### TAG assay

Female flies were collected and divided into groups of 5, 3 days post eclosion. Each group was weighed on a Thermo microbalance to an accuracy of .01 mg. Weighed females were added to a sterile 2mL screw-cap tube containing sterile 1.4mm ceramic beads and 125 µL of TET buffer (10mM Tris, 1mM EDTA, pH 8.0, 0.1% (v/v) Triton-X-100). Samples were homogenised for 30s at max speed. After homogenisation, samples were incubated at 72°C for 15 min in a dry bath to inactivate lipases and other enzymes. Then samples were centrifuged at max speed for 5 min at 4°C. From each sample 3 µL of the supernatant was added to a clear, flat bottom 96 well plate. Standard curves were generated using a glycerol stock ranging from 1-0 ug/ul. Triglyceride levels were assayed by the method of Fossati and Prencipe(Fossati and Prencipe, 1982), with absorbance measured at 540 nm in a Thermo MultiSkan FC plate reader.

### RNA extraction

RNA was extracted from whole flies by the addition of 10 female flies per sample to sterile 2 mL screw cap tubes containing 600 µL of TRIzol (Ambion by Life technologies) and homogenised in a Ribolyzer and running at 6.5 (max) for 10 seconds. 200 µL chloroform was added, briefly vortexed, and incubated at RT for 3 min. Samples were centrifuged 12,000rpm for 15m at 4°C, 250µl of isopropanol was added to supernatants, vortexed, and incubated at RT for 10 minutes. Samples were then centrifuged at 12,000rpm for 10m at 4°C. The liquid was removed and discarded, while the pellet containing the RNA was washed with 1mL of 75% ethanol. Samples were centrifuged at 12,000 rpm for 10m at 4°C. The liquid was removed without disturbing the pellet, and the tubes were dried at RT with caps opened. Pellets were reconstituted in DEPC water (10µl/fly) and stored at −80°C.

### 16s amplicon sequencing

DNA was extracted from whole guts, dissected from 3 day-old wild-type stock females (i.e. conventionally-reared and bearing a complete microbiota, 5 per sample, 10 samples total) after 1 minute surface-sterilisation in 70% EtOH. Guts were deposited immediately into TRIzol and homogenised in a ribolyser. 300µl chloroform was added and inverted followed by 3 mins incubation at room temperature, 15 minutes centrifugation at 12000g. Aqueous layer was removed before 500µl back extraction buffer (4M Guanidine thiocyanate, 50 mM sodium citrate, 1M Tris base) was added and mixed by inverting. Samples were spun again at 12000g for 30 minutes and aqueous phase transferred to new tubes with 400 µl isopropanol. Samples were mixed by inversion, centrifuged 15 minutes at 12000g. Pellets were washed in 70% ethanol, dried, and resuspended in 50 µl TE buffer.

Library prep was performed by the University of Glasgow MVLS Shared Research Facility from 12.5ng input DNA per sample, with an initial PCR using Accustart PCRToughMix and 0.5µl 10mM primers against 16s V3/V4 region (FWD TCGTCGGCAGCGTCAGATGTGTATAAGAGACAGCCTACGGGNGGCWGCAG, REV GTCTCGTGGGCTCGGAGATGTGTATAAGAGACAGGACTACHVGGGTATCTAATCC), using 25 cycles at an annealing temperature of 60°C, followed by barcoding with Illumina Nextera XT 1 primers, cleanup with 0.9X Spri Select beads (Beckman Coulter), and washing in 80% EtOH. Libraries were quantified by Qubit HS DNA assay and size distributions checked on an Agilent Bioanalyser. Pooled libraries were sequenced on an Illumina MiSeq in paired end mode (300bp) using a 600 cycle cartridge kit, 5% PhiX, to an average depth of 170K reads per sample.

### RT-qPCR

RNA samples were quantified by spectrophotometry using a Nanodrop 1000 UV-Vis spectrophotometer (ThermoScientific). First strand synthesis of cDNA was performed using the SuperScriptTM Reverse Transcriptase (RT) kit (Invitrogen), following the manufacturer’s protocol (using 2µg of RNA). The PCR profile was as follows: 95°C for 2 minutes; 94°C denaturation for 30 seconds, 60°C annealing for 30 seconds, 72°C extension for 30 seconds each for 40 cycles, with a final extension at 72°C for 5 minutes. The amplicon was migrated through a 1% (w/v) agarose gel. To test for relative gene expression levels, RT-qPCR was performed. Each well contained 2 μl cDNA, 1.5 μl 10 mM forward primer, 1.5 μl 10 mM reverse primer, 3 μl water, 10 μl SYBR green PCR master mix, and 2 ul ROX (Qiagen, UK). The following PCR profile was used: 95°C for 5 minutes; 94°C for 30 seconds, 60°C for 30 seconds, 72°C for 30 seconds each for 40 cycles, and a final extension at 72°C for 5 minutes. Mean cycle threshold (Ct) values for duplicates of each of the studied genes were derived by subtracting the combined mean Ct value of constitutively expressed housekeeping genes *alcohol dehydrogenase* (*ADH*) or *tubulin*. Primers shown in Table 4.

**Table 4.**
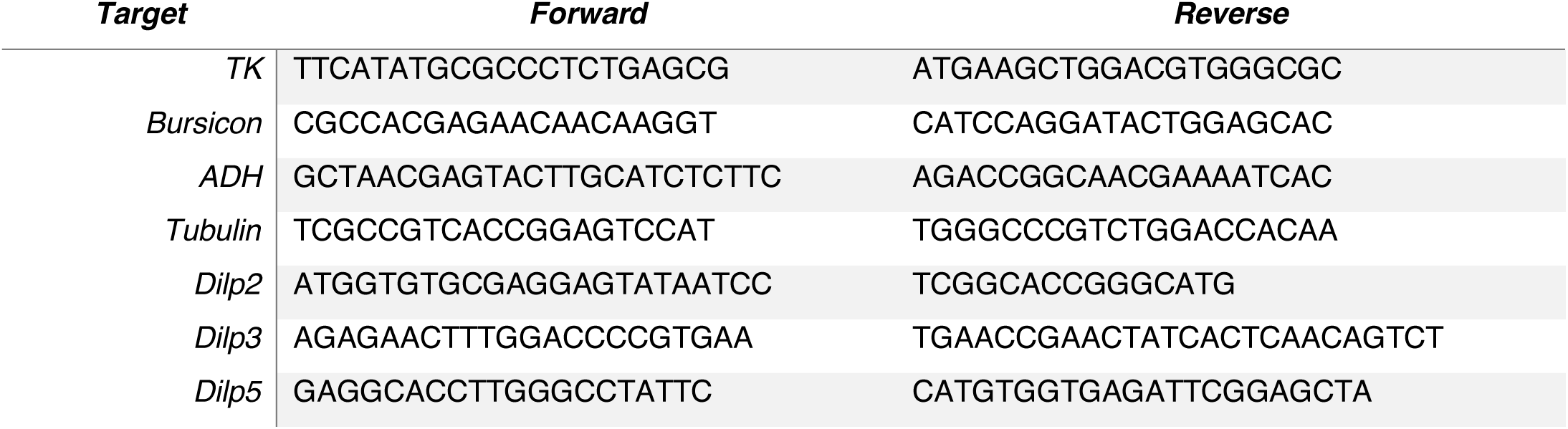
Primers for RT-qPCR.

### Antibody staining and imaging

Whole guts were extracted from 10-day old female flies (axenic, gnotobiotic, and axenic on 1µM TSA), and fixed in 4% paraformaldehyde for 30m in a 12 well plate. Guts were then incubated in 1 mL of PBST (1X PBS with 0.1% triton X-100) overnight with slight shaking at 4°C. PBST was carefully aspirated from sample wells, and guts were then washed with 1 mL of Blocking buffer (10% 1X PBS, 1% triton X-100, 2.5% Normal Goat Serum) at 4°C overnight. Blocking buffer was removed, and 0.5 mL of acetyl-lysine antibody (Cell Signaling Technologies 9441) diluted 1/800 in blocking buffer was added to the wells, and incubated overnight at 4°C with slight shaking. Guts were then washed 3X with PBST for 10 minutes, followed by a 1 hr RT PBST wash with slight shaking. 0.5 mL of secondary antibody secondary antibodies diluted 1/200 in PBST was added to the guts, and incubated overnight at 4°C. Guts were washed 3X in PBST for 10 minutes. Guts were then mounted in Vectashield for imaging.

Corpora cardiaca were dissected, fixed and washed as above, stained with anti-Akh (gift from Kenneth Halberg) diluted 1:1000, and stained with secondary antibodies as above.

### Confocal microscopy

Imaging was performed on a Zeiss LSM 880 confocal microscope and processed using Zeiss Zen Blue (v3.8) and NIH ImageJ software 1.53a prior to transfer to Adobe Photoshop and Illustrator (Creative Cloud 2025; CA, USA) for final presentation.

## Data analysis

All enumerated data were plotted and analysed in R 4.4.2. (Team, 2024). Kaplan-Meier curves were plotted with survminer::ggsurvplot_facet. Other quantitative data were plotted with ggplot2. Survival data were analysed with CoxPH models (survival::coxph). Fecundity data were analysed with negative binomial models (lme4::glmer.nb). All other data were analysed with stats::lm (after logit transformation for proportion flies feeding). Post-hoc tests were performed with emmeans::joint_tests and emmeans::pairs, coefficients were extracted from emmeans::emmip. Survival indices were generated by applying emmeans::emmip to coxph models, multiplied by −1, such that higher values corresponded to longer lifespan.

Published transcriptome data (Bost et al., 2018) were reanalysed by FastQC and quantified with Salmon. The previous publication by Bost et al(Bost et al., 2018) presented bulk RNAseq on midguts of flies reared under axenic conditions or with a 5-species gnotobiotic microbiota comprising *Acetobacter* and *Levilactobacillus* species. The original analysis presented excluded expression of ∼70% of genes in the fly genome, and hormones were largely excluded. We reanalysed the raw data using Salmon (Patro et al., 2017) to quantify expression, which quantified expression of 10,290 genes, including 14 of the 15 enteroendocrine hormone genes (except *bursicon*). We tested for differential expression using DESeq2, identifying 470 differentially expressed genes (FDR ≤ 0.01, ≥2-fold difference in mean expression). These 470 genes included four peptide hormones: *Tachykinin (Tk), Allostatin C (AstC), Ion Transport Peptide (ITP), and Neuropeptide F (NPF)*. Since *Burs* was not quantified by the Salmon algorithm, we tested independently whether its expression was modulated by microbiota by extracting RNA from midguts of CR and axenic flies.

## Acknowledgments

We thank Julia Cordero for helpful discussion, reagents and training; Nathan Woodling and Colin Selman for comments on the manuscript; and Kim Rewitz, Kenneth Halberg, Jean-René Martin and Peter Newell for reagents. The authors gratefully acknowledge the Cellular Analysis Facility at the University of Glasgow – particularly David McGuinness – for support and assistance in this work. This work was funded by a UKRI Future Leaders Fellowship (MR/S033939/1 and MR/Y019660/1, AD), BBSRC (BB/W510658/1, DRS and AD), a University of Glasgow Lord Kelvin Adam Smith Fellowship (AD), a University of Glasgow Lord Kelvin Adam Smith Scholarship (RI), and funds from the University of Glasgow Wellcome Trust Institutional Strategic Support award (RI and AD).

## Supplementary figure legends

**Figure S1.**
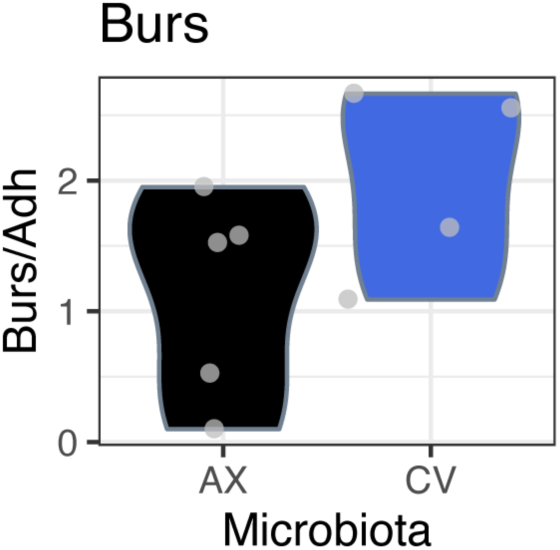
Microbiota do not alter intestinal *Burs* expression. Expression was quantified by RT-qPCR (using cDNA generated from midgut to complement RNA-seq data) because transcripts were not quantified from RNA-seq data (Figure 1). No significant difference in expression was detected (t=-1.65, p=0.15). AX=axenic, CV=conventionally-reared. Expression was quantified relative to *Adh*, as a housekeeping gene that RNAseq data showed to not be influenced by microbiota (Figure S2)

**Figure S2.**
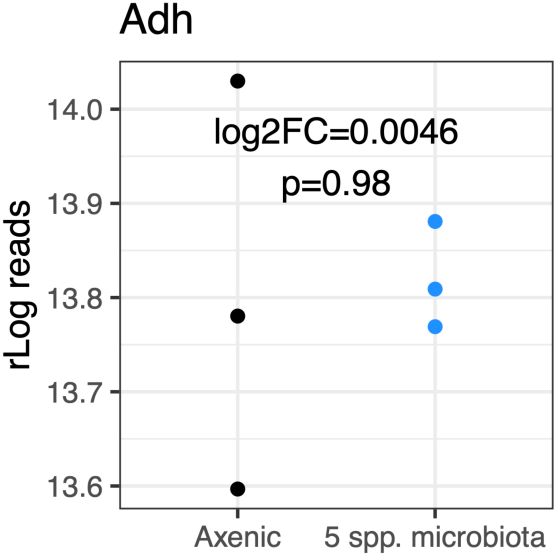
Microbiota do not alter intestinal *Adh* expression. Expression was quantified from public RNAseq data (Bost et al., 2018). Statistics show log2 fold-change and p-value (unadjusted) from DESeq2 analysis of reads quantified with Salmon.

**Figure S3.**
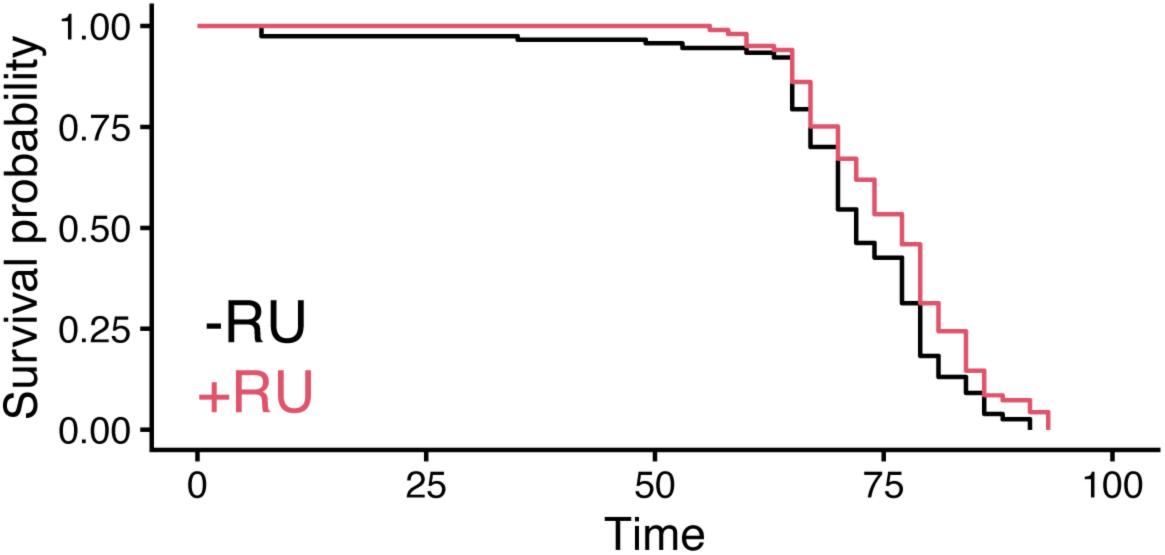
Lifespan extension in conventional flies by induction of an independent *Tk^RNAi^*. To validate result show in Figure 2A a distinct construct (Vienna *Drosophila* Stock Center #25800) was expressed under control of *DaGS.* Log-rank test p<0.01. -RU condition 108 deaths, 43 censors; +RU condition 124 deaths, 26 censors.

**Figure S4.**
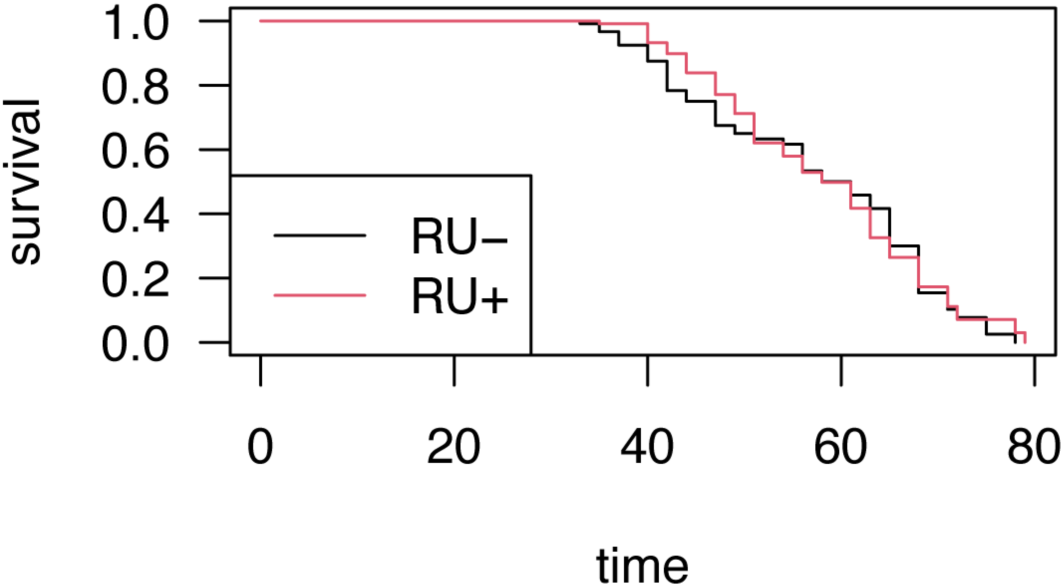
Lifespan extension by feeding RU to CR *DaGS;UAS-Tk^RNAi^* flies is not attributable to off-target effects of RU or GeneSwitch activation. Kaplan-Meier plot shows impact of feeding RU to *DaGS/+* flies reared with complete microbiota. No effect of RU was detected (Cox Proportional Hazards p=0.5).

**S5.**
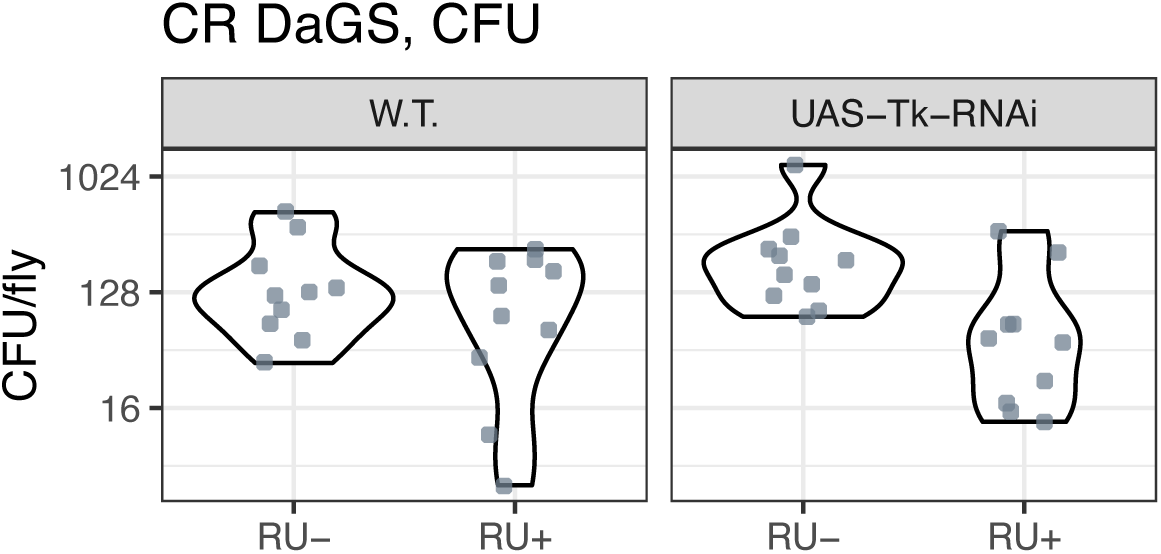
CFU. *TkRNAi* does not affect bacterial load in CR flies. Violin plots show CFUs of conventionally-reared female adults of the indicated genotypes (all with *DaGS*), after one week of feeding on RU or vehicle control. All conditions n=10. No effects were detected of presence/absence of *UAS-Tk-RNAi* (ANOVA of log CFU counts, p=0.32), RU (p=0.23), nor genotype:RU interaction (p=0.25).

**Figure S6.**
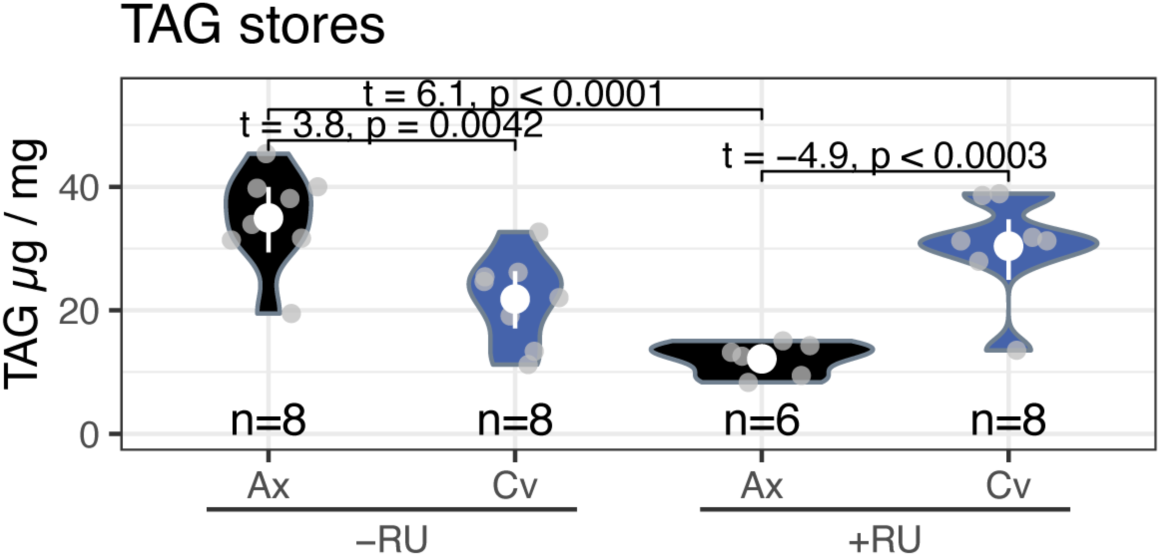
Interactive effect of *TkRNAi* and microbiota on TAG levels confirmed with independent RNAi construct. The experiment presented in Figure 2E (which used construct V330743) was repeated with an independent construct (V103668). The same result was replicated. Violin plots show density of datapoints, mean±CI95 given in white. ANOVA (Type 3 tests), microbiota:RU F_1,26_=37.86, p=1.66e-06. Statistics on brackets are from *post-hoc* EMM analyses of linear model, when significant differences were detected.

**Figure S7.**
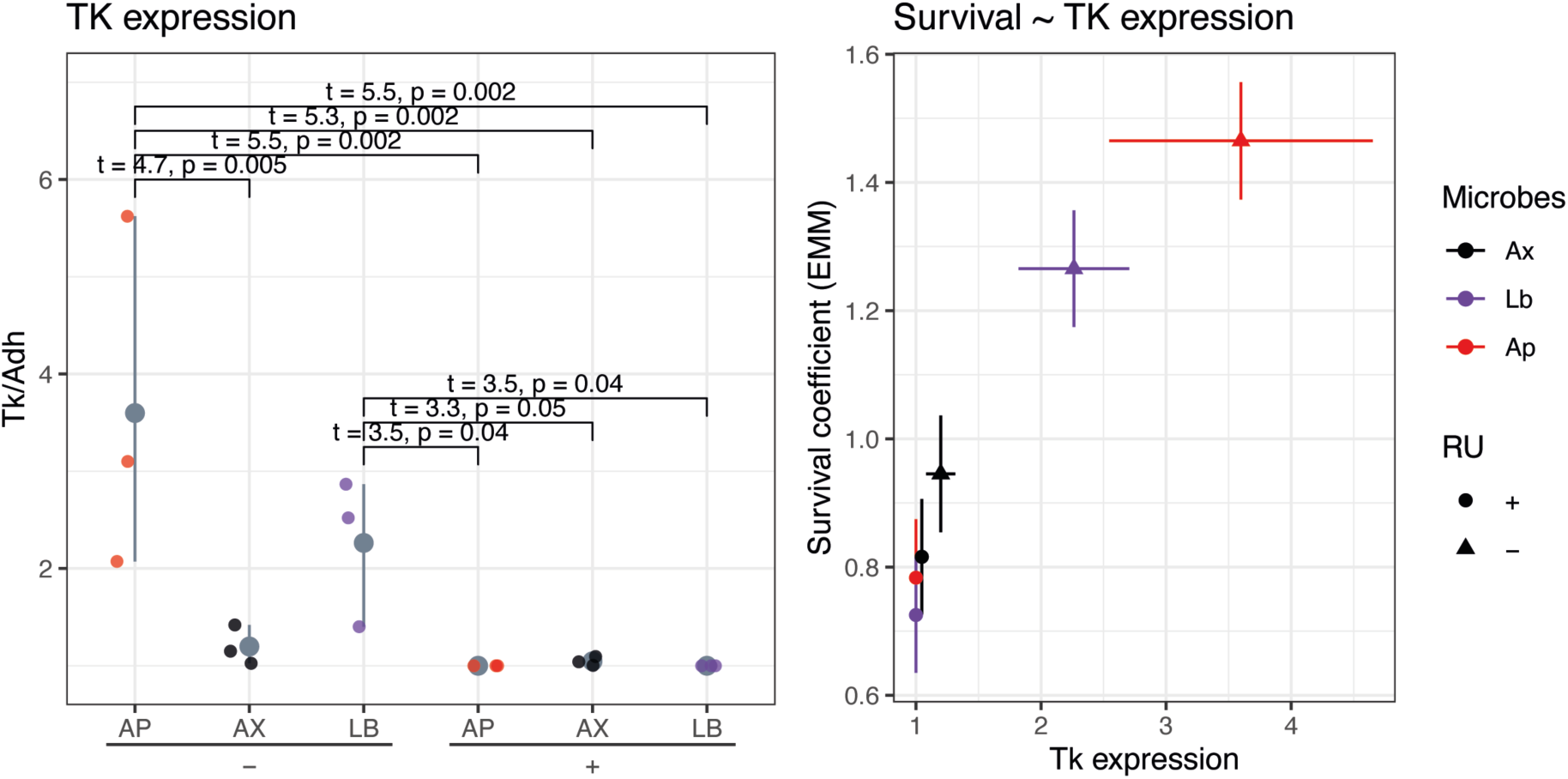
Modulation of *Tk* expression by microbiota correlates lifespan outcome. **A.** RT-qPCR indicates elevated *Tk* expression in *Ap-*flies above levels in axenics and *Lb-*flies, and above levels in all conditions expressing *Tk^RNAi^*. Expression in *Lb-*flies was significantly greater than in conditions expressing *Tk^RNAi^,* but not significantly greater than axenics without *Tk^RNAi^* expression. Statistics on brackets are from *post-hoc* EMM analyses of linear model, when significant differences were detected. **B.** *Tk* expression differences across microbiota and *Tk^RNAi^* conditions correlate lifespan differences. *Tk* expression from panel A is plotting against survival coefficients of lifespan experiment shown in Figure 3c-d.

**Figure S8.**
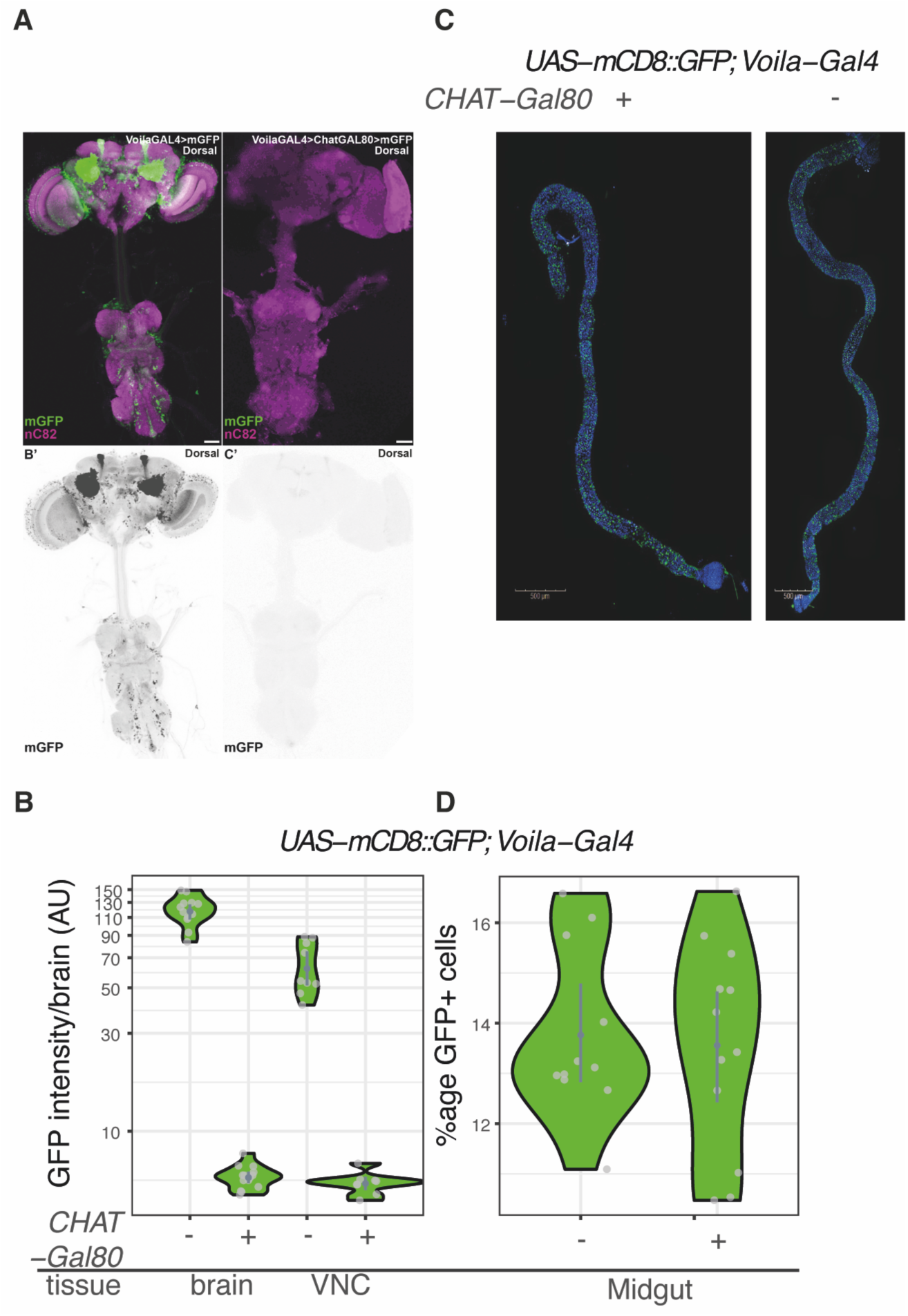
Intersectional Gal80 strategy to direct Gal4 activity to enteroendocrine cells, sparing CNS. As per Medina et al (Medina et al., 2024), we tested the capacity of Gal80 expressed under control of the *ChAT* promoter to repress Gal4 expressed under control of the *Voila* promoter, assayed by expression of *UAS-mCD8::GFP.* **A.** Representative confocal micrographs of CNS, and **B.** Quantification of Voila>GFP intensity in brain and CNS, in presence/absence of *Voila Gal80*. **C.** Representative confocal micrographs of midgut (region 4). **D.** Number of GFP+ cells/ROI in presence/absence of *Voila Gal80*.

**Figure S9.**
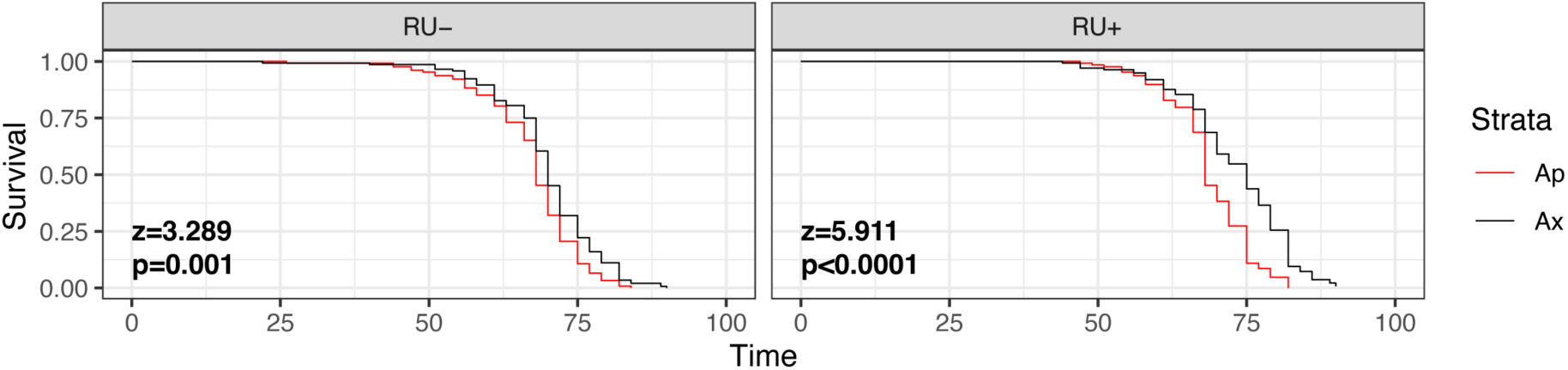
*TkR99D^RNAi^* induction in enterocytes does not block impact of *A. pomorum* on lifespan. Expression was driven with *Mex1-GeneSwitch*. Statistics in panel from Cox proportional hazards model, reporting post-hoc (EMM) analysis for effect of RU per microbiota condition.

**Figure S10.**
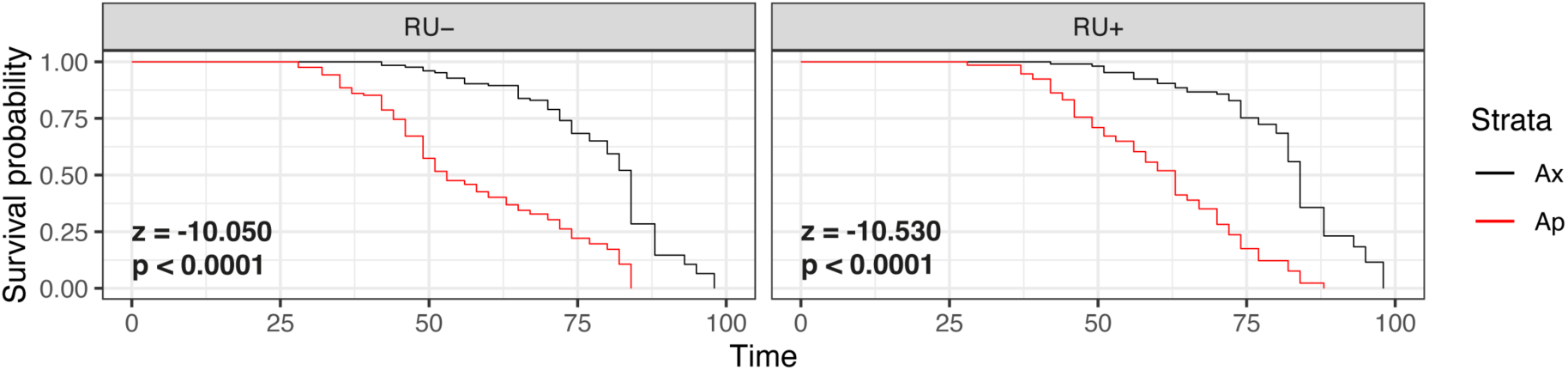
*AkhR^RNAi^* induction in fat body (and gut) does not block impact of *A. pomorum* on lifespan. Expression was driven with *S106-GeneSwitch*. Statistics in panel from Cox proportional hazards model, reporting post-hoc (EMM) analysis for effect of RU per microbiota condition.

